# Gene therapy for targeting a prenatally enriched potassium channel associated with severe childhood epilepsy and premature death

**DOI:** 10.1101/2024.10.24.620125

**Authors:** Sean R. Golinski, Karla Soriano, Alex C. Briegel, Madeline C. Burke, Timothy W. Yu, Tojo Nakayama, Ruilong Hu, Richard S. Smith

## Abstract

Dysfunction of the sodium-activated potassium channel K_Na_1.1 (encoded by KCNT1) is associated with a severe condition characterized by frequent seizures (up to hundreds per day) and is often fatal by age three years. We defined the early developmental onset of K_Na_1.1 channels in prenatal and neonatal brain tissue, establishing a timeline for pathophysiology and a window for therapeutic intervention. Using patch-clamp electrophysiology, we observed age-dependent increases in K_Na_1.1 K^+^conductance. In neurons derived from a child with a gain-of-function KCNT1 pathogenic variant (p.R474H), we detected abnormal excitability and action potential afterhyperpolarization kinetics. In a clinical trial, two individuals with the p.R474H variant showed dramatic reductions in seizure occurrence and severity with a first-in-human antisense oligonucleotide (ASO) RNA therapy. ASO-treated p.R474H neurons in vitro exhibited normalized spiking and burst properties. Finally, we demonstrated the feasibility of ASO knockdown of K_Na_1.1 in midgestation human neurons, suggesting potential for early therapeutic intervention before the onset of epileptic encephalopathy.

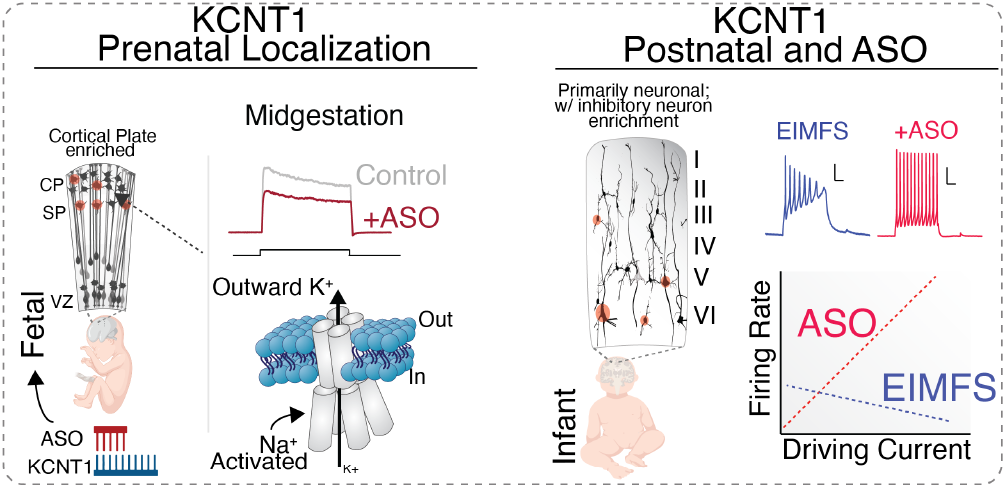

## Introduction

RNA therapeutics offer exciting new clinical opportunities for treatment-resistant neurodevelopmental disorders, including treatments to modify the RNA expression of difficult-to-treat proteins using targeted antisense oligonucleotides (ASOs), which was first successfully demonstrated to treat spinal muscular atrophy (SMA)^1^. Additional investigational ASO therapies for orphan neurogenetic diseases have been clinically successful^2,3^; for example, ASO treatment was successful in two children with drug-resistant KCNT1-associated epilepsy of infancy with migrating focal seizures (EIMFS)^4^, a severe condition in which half of the patients do not survive past the age of 3 years^5^. Following ASO treatment, these KCNT1 patients showed a marked reduction in seizure frequency and intensity4, offering clinical promise for potential therapeutic interventions prior to encephalopathy, loss of developmental milestones, and death^5^.

KCNT1 early infantile epileptic encephalopathy (EIEE14) is a severe epilepsy syndrome affecting newborns^6^, resulting from gain-of-function (GoF) de novo variants of KCNT1 that increase outward K^+^ currents^7,8^. In the central nervous system, KCNT1 encodes an brain enriched sodium-activated potassium channel (K_Na_1.1)^9^ that supports critical neuronal functions in the postnatal brain, such as burst firing and action potential (AP) kinetics^10^. Functional characterization of a recurrent EIEE14 variant (p.R474H) in KCNT1 demonstrated a robust GoF channel activity phenotype that is resistant to treatment in newborns^7,8,11^. While ion channel dysfunction is known to disrupt human prenatal processes, resulting in severe neurodevelopmental disease^12^, the contribution of K_Na_1.1 to normal brain development and the potential for therapeutic targeting before birth is unknown.

To address this gap, we utilize a novel RNase H-dependent gapmer ASO, recently used to reduce seizures in two individuals affected with KCNT1-associated EIMFS^4^, to define the mechanistic recovery of action potential pathophysiology in patient-derived neurons. Furthermore, we profiled the developmental emergence of K_Na_1.1 currents in primary human neurons and demonstrated that KCNT1 can be effectively knocked down using an ASO as early as 16 weeks gestation. The emergence of functional K_Na_1.1 during fetal development and its contribution to neuronal excitability in utero suggest that GoF variants in K_Na_1.1 contributes to disease pathophysiology prenatally. This finding suggests the possibility of improving clinical outcomes by applying ASO treatment prenatally or in early infancy.

## Results

### EIMFS neurons treated with ASO generate pronounced afterhyperpolarizations attenuated by blockers of calcium-activated K^+^ channels

The pathogenic variant KCNT1-p.R474H is associated with EIMFS in children with dozens to hundreds of treatment-resistant multifocal seizures per day that can be reduced in number and severity by a targeted ASO^4^. To assess the effect of the ASO on p.R474H neurons, we obtained voltage-clamp recordings of cortical excitatory neurons generated via NGN2 directed differentiation from human induced pluripotent stem cells (hiPSCs) from an EIMFS individual with this variant. After maturing p.R474H neurons in culture for 14 days (DIV 14), we treated the neurons for 14 days (until DIV 28) with a KCNT1-targeting ASO (referred to as ASO), or a control non-hybridizing sequence ASO with related thiophosphate chemistry (referred to as NT-ASO). We demonstrated knockdown of KCNT1 by ASO in p.R474H neurons using quantitative reverse transcription polymerase chain reaction, whereas NT-ASO did not affect KCNT1 RNA levels (Figure S1B), confirming previous work4. Voltage-clamp recordings of KCNT1-p.R474H neurons showed an increased steady-state outward K^+^ current (consistent with GoF) compared with control neurons (PGP-1 cell line, Figure 1B). ASO treatment of KCNT1-p.R474H neurons reduced outward K+ current (vehicle vs. ASO: p = 0.002), whereas NT-ASO did not reduce K^+^ current (vehicle vs. NT-ASO: p = 0.7; Figure 1B). Critically, p.R474H neurons treated with ASO, NT-ASO, or vehicle displayed comparable levels of maximum inward sodium current density (Figure S1D, p = 0.988); therefore, the level of sodium activating the K^+^current is unlikely to underlie the ASO effects. To analyze potential ASO-induced toxicity or changes to neuronal morphology among the ASO- and NT-ASO-treated neurons, we performed post-hoc morphological analysis (Figure S1C). We did not detect any differences in the ramification index, soma area, or length of the longest neurite of the neurons among the three conditions (Figure S1B, p > 0.05), suggesting that the 14-day ASO and NT-ASO treatments do not alter complex neuronal morphology.

**Fig. 1.**
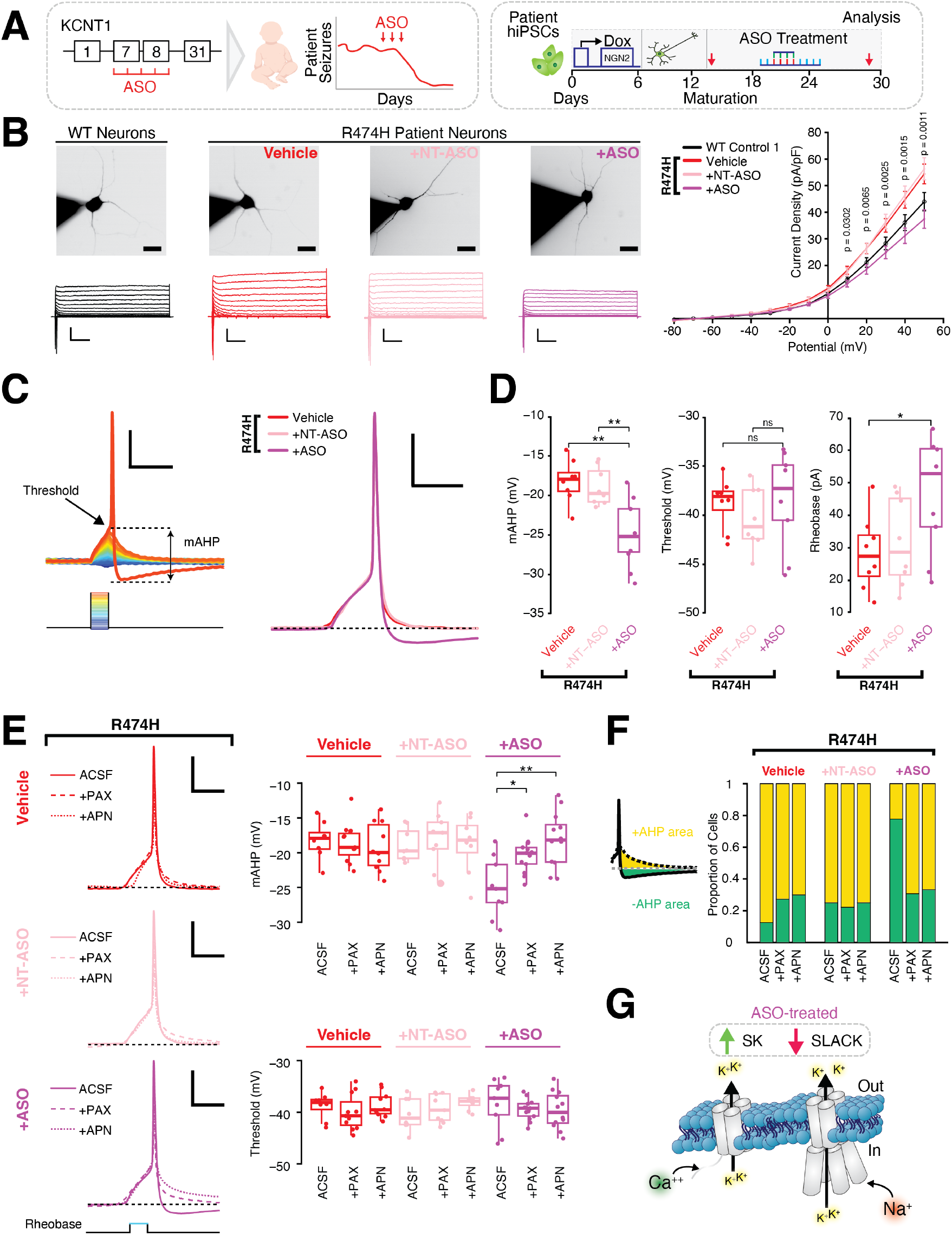
A. Left, ASO design (5-10-5 gapmer) targeting exons 7 and 8 of KCNT1, and a representative time graph of seizure reduction in a KCNT1-p.R474H patient following ASO treatment4. Right, schematic of NGN2 neuronal differentiation of iPSCs from a patient with the KCNT1-p.R474H variant. B. Top, representative epifluorescence images of NGN2 neurons loaded with Alexa-Fluor 488 dye (1 μM) during patch recordings (scale bar: 10 μm). Bottom, voltage clamp traces evoked by step activation routine from -80 to +50 mV at 10 mV increments (scale bars: 1 nA/100 ms). Right, IV plot showing KCNT1-p.R474H NGN2 neurons have increased GoF steady-state outward K+ current densities sensitive to ASO knockdown (p = 0.0018 at 50 mV) and unaffected by a control NT-ASO (p = 0.74 at 50 mV). P-values shown are between neurons treated with NT-ASO and ASO. C. Left, Representative AP from NGN2 neuron and sequential depolarizations leading to first firing using a rheobase routine (40 ms current injections from -5 pA to +80 pA at 2 pA increments) with Vm = -60 mV. Right, average first AP elicited across conditions, showing significantly larger mAHP in cells treated with ASO (scale bars: 20 mV/100 ms). Single cell Aps overlayed with average AP in S1. D. Left, KCNT1-p.R474H neurons display larger mAHPs following ASO treatment (p = 0.013). Middle, average thresholds are comparable across all conditions (n.s., p > 0.8). Right, KCNT1-p.R474H neurons treated with ASO require significantly more stimulation to achieve the first AP, rheobase current (p = 0.036). E. Left, Average first AP elicited across conditions following bath perfusions of BK blocker paxilline and SK blocker apamin (PAX 500 nM and APN 50 nM, respectively), showing mAHP reduction in ASO-treated neurons. Right, only ASO-treated KCNT1-p.R474H neurons were sensitive to mAHP reduction following bath application of PAX and APN (p = 0.044, p = 0.002, respectively). Threshold values are unaffected by PAX and APN. F. Proportion of cells that netted a negative mAHP area (as measured by taking the area with respect to the holding potential of -60 mV). A significant decrease in the proportion of cells netting a negative mAHP area in ASO-treated cells is observed following PAX and APN perfusion. G. Schematic depicting hypothesized upregulation of SK channels in response to ASO-mediated KCNT1 knockdown. For all ASO treatments of NGN2 neurons (14 days), the initial dose was 10 μM (at DIV 14), with a second 5 μM maintenance dose at DIV 21. * p <0.05, ** P <0.01. See Table S1 for statistics.

Quantification of AP rheobase, a value that describes the minimum input current (in pA) necessary to evoke an AP (Vm holding = –60mV), revealed paradoxically increased rheobase in KCNT1-p.R474H neurons treated with ASO (vehicle: 28.3 ± 4.0 pA, n = 8; ASO: 46.0 ± 5.6 pA, n = 9, p = 0.025; Figures 1C, D), yet KCNT1-p.R474H neurons treated with NT-ASO exhibited a comparable rheobase to that of vehicle neurons (NT-ASO: 31.73 ± 4.79 pA, n = 8, p = 0.59; Figure 1D, Table S1). Across KCNT1-p.R474H neurons treated with vehicle or ASO, we did not detect changes in cell capacitance (Vehicle vs. ASO: p = 0.565) or input resistance (Vehicle vs. ASO: p = 0.218, Table S1).

Next, in ASO-treated KCNT1-p.R474H neurons, we analyzed medium afterhyperpolarization (mAHP), a key property of AP recovery carried by K+ conductances. During current clamp, we applied the minimal current stimulation needed to elicit a single AP with a reliable mAHP. We observed an increased mAHP amplitude in ASO-treated KCNT1-p.R474H neurons compared with NT-ASO- and vehicle-treated neurons (Vehicle: –18.2 ± 0.9 mV, n = 8; ASO: –25.1 ± 1.4 mV, n = 9; p = 0.001; Figure 1D, Table S1). As mAHP is measured with reference to the neuron AP threshold, we confirmed that the threshold values were comparable across conditions (vehicle: –38.76 ± 0.91 mV, n = 8; ASO: –38.34 ± 1.595 mV, n = 9, p = 0.831; Figure 1D). In a control hiPSC line from a healthy individual (PGP1, unrelated to proband), we also observed that ASO-treated neurons result in increased mAHP (PGP1: -20.07 ± 1.19 mV, n = 9; PGP1+ASO: -24.93 ± 1.27 mV, n = 10, p = 0.013; Figure S1D, Table S1).

To isolate the ionic conductance underlying the ASO increase in mAHP amplitude, we perfused selective antagonists of the voltageand Ca^2+^-activated large-conductance (BK) channel (paxilline, PAX, 500 nM) and of the Ca^2+^-activated small-conductance (SK) potassium channel (K_Ca_2) (apamin, APN, 50 nM). The increased mAHP observed in ASO-treated p.R474H neurons decreased significantly with PAX and, to a larger degree, with APN (Figure 1E; baseline (ASO): –25.05 ± 1.432 mV, n = 9; ASO + PAX: –20.36 ± 0.710 mV, n = 13, p = 0.011; ASO + APN: –18.107 ± 1.132 mV, n = 12, p = 0.002, See Table S1), yet we did not observe any changes with PAX or APN in vehicle-or NT-ASO-treated cells (Figure 1E, Table S1). Similarly, the BK and SK channel antagonists (PAX and APN) did not impact thresholds for the vehicle, NT-ASO, or ASO conditions (Figure 1E, Table S1). Furthermore, the voltage-gated Ca^2+^ channel blocker nifedipine (100 *μ*M) did not affect the mAHP amplitude in ASO-treated KCNT1-p.R474H neurons (Figure S1H, p = 0.28).

In addition to the mAHP amplitude, we analyzed the proportion of cells that had a net negative mAHP area under the resting membrane potential (–60 mV). Our results demonstrated that ASO-treated p.R474H neurons perfused with PAX or APN displayed a reduced percentage of cells that would hyperpolarize below the RMP, resulting in more cells with a positive net mAHP area (Figure 1F).

### ASO-treated KCNT1-p.R474H neurons display an improved dynamic range of excitability

KNa1.1 regulates neuronal burst rate and AP kinetics^10^, including AHP kinetics and maintenance of rhythmic burst recurrence during sustained depolarization^13^. To characterize the bursting and excitability properties of EIMFS KCNT1-p.R474H neurons treated with ASO, we injected a wide range of currents (–10 to 180 pA in 10-pA steps) to generate sustained depolarizations and APs (Figure 2A). In ASO-treated p.R474H neurons, we observed a bidirectional effect in the input–output firing curves compared with vehicle- and NT-ASO-treated neurons. At low current injections, ASO treatment reduced the excitability (at 40 pA, vehicle: 10.55 ± 0.86 APs, n = 10; ASO: 5.33 ± 1.85 APs, n = 12, p = 0.023; Figure 2C); however, at higher current injections, ASO-treated cells fired at higher frequencies than vehicle- and NT-ASO-treated cells (at 180 pA, vehicle: 5.00 ± 0.58 APs, n = 10; ASO: 18.22 ± 2.17 APs, n = 12, p = 0.006; Figure 2C). ASO treatment also improved the AP amplitude run-down between spikes, yet did not significantly alter the spike frequency adaptation (Figure 2D), demonstrating a recovery of AP kinetics during burst trains.

**Fig. 2.**
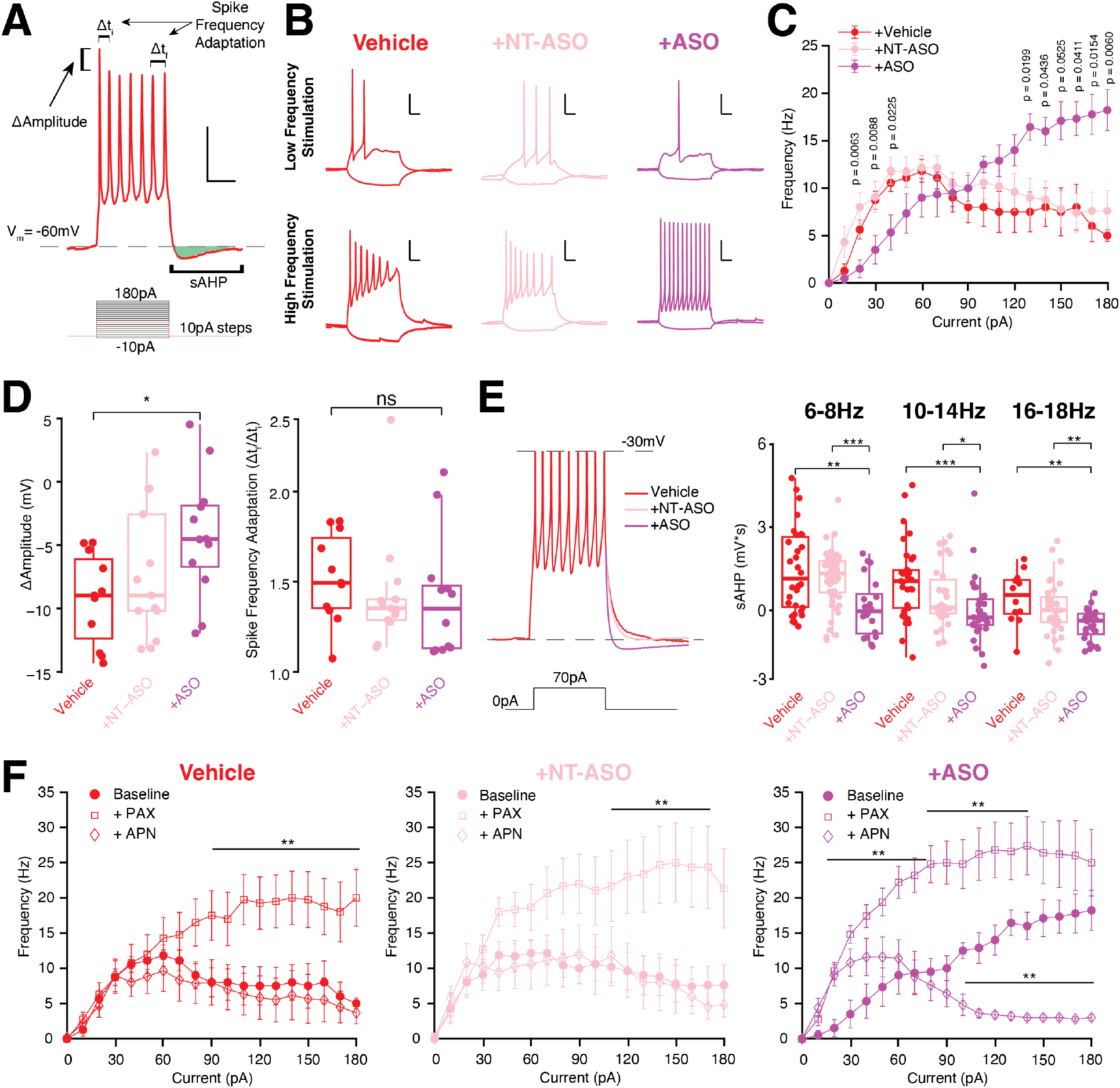
A. Representative AP burst of a KCNT1-p.R474H neuron at Vm= -60 mV in response to prolonged current injection with analysis parameters defined (−10 pA–180 pA, 10 pA steps, 500 ms). B. Representative AP response profiles of KCNT1-p.R474H neurons following 14-day ASO treatment protocol (vehicle, ASO, and NT-ASO) at low- and high-frequency stimulation (20 pA and 120 pA, respectively). C. Frequency plotted as a function of input current reveals that ASO-treated KCNT1-p.R474H neurons fire at significantly higher frequencies at large current injections compared to vehicle or NT-ASO (p=0.006, p=0.008, respectively). D. KCNT1-p.R474H neurons displayed a marked reduction of first to second AP amplitudes (Δ Amplitude), which was recovered by the ASO treatment, without any significant effect on spike frequency adaptation. E. Average sAHP elicited following spiking bursts for all traces between 70 pA and 90 pA (where spiking frequencies were most alike across conditions). ASO-treated neurons display increased (more negative) sAHP area. Right, frequency-dependent sAHP area is increased in ASO-treated neurons over a range of frequencies, including at low (6–8 Hz), medium (10–14 Hz), and high (16–18 Hz) frequencies. F. Input–output excitability curves of vehicle-, NT-ASO-, and ASO-treated KCNT1-p.R474H neurons following bath perfusion of PAX (500 nM) and APN (50 nM) demonstrate that ASO-treated neurons rely on SK channels for maintaining firing at higher current injections, likely by enhancing AHP and delaying depolarization block at high current injections. BK channel blockers enhanced firing in all conditions. For all ASO treatments of NGN2 neurons (14 days), the initial dose was 10 μM (at DIV 14), with a second 5 μM maintenance dose at DIV 21. Data are presented as mean±SEM. * p <0.05, ** P <0.01, *** P <0.001. See Table S2 for statistics.

Because ASO-treated KCNT1-p.R474H neurons displayed improved run-down and a dynamic range of input–output AP spiking (Figures 2C, D), we next explored slow AHP (sAHP). sAHPs are frequency-dependent and critical to neuron firing, as they are generated following trains of synaptic input and burst AP firing, and are carried by Ca2+-dependent K+ channels14. ASO-treated KCNT1-p.R474H neurons displayed larger sAHPs than vehicle- and NT-ASO-treated neurons across a range of firing potentials (p = 0.002 at 6–8 Hz, p < 0.001 at 10–14 Hz, and p = 0.002 at 16–18 Hz; Figure 2E). Next, we pharmacologically isolated ionic conductances underlying ASO improvement in AP firing dynamics using BK and SK channel agonists (PAX, 500 nM, APN, 50 nM, respectively). Intriguingly, whereas the BK antagonist (PAX) increased high-frequency firing across all three conditions (vehicle, NT-ASO, and ASO), the SK blocker (APN) reduced high-frequency AP firing only in the ASO-treated KCNT1-p.R474H condition (Figure 2F, Table S2). Moreover, the KCNQ2 channel antagonist XE991 did not reduce AP excitability in ASO-treated neurons, but instead increased high-frequency firing (ASO vs. ASO+XE991: p < 0.05 at all current injections; Figure S2C). These highly selective pharmacological treatments indicate that ASO-treated KCNT1-p.R474H neurons likely rely on Ca2+-activated SK channels (KCa2) to support the observed high-frequency AP firing.

### KCNT1 is expressed in mid-gestation human cortical plate neurons

Early diagnosis and intervention for epilepsy disorders improve clinical outcomes, yet drug-resistant infantile epilepsies have poor outcomes^15^. KCNT1-associated EIMFS is typically observed within 6 months after birth^16,17^, suggesting that KCNT1 pathology might occur before this period. Therefore, we sought to define a cell-type-specific prenatal profile of KCNT1 leading up to the neonatal period. Analysis of bulk cortical human transcriptome data from the Allen Brain Atlas18 from 12 weeks post-conception (PCW) to adulthood (40 years) indicated that KCNT1 RNA expression is first observed mid-gestation (13–28 PCW) and persists into adulthood (Figure 3A). This developmental emergence of KCNT1 expression coincides with several overlapping processes, including neuronal migration and differentiation, which can be disrupted by ion channel dysfunction12. To investigate KCNT1 cell-type distribution and potential spatiotemporal-dependent pathology, we performed spatial RNA profiling in the human fetal neocortex during the mid-gestation KCNT1 onset window (16–21 PCW). We applied multiplexed fluorescent in situ hybridization analysis of KCNT1 co-expression within developmental cell types along a neocortical column and found that KCNT1 mRNA was not detected in the progenitor germinal zones, including within radial glia or immediate progenitors (EOMES- and Vimentin-positive cells) of the ventricular and intermediate zones (Figure 3B). Similarly, we did not observe KCNT1 expression in progenitors of the outer subventricular zone or within the primate-enriched outer radial glial cells (HOPX+) (Figure 3B). The primary KCNT1 mid-gestational expression occurs within postmitotic neurons (RBFOX3+) in the early cortical plate (Figure 3). We did not observe KCNT1 expression within immature migrating neurons (Figure 3B) or subplate neurons (Figure S3A), in contrast to the cell-type expression patterns observed for other early epilepsy-related ion channel diseases^19^. Neuron-subtype-specific markers reveal KCNT1 expression within several excitatory cell types in the cortical plate, including deep-layer neuron markers TBR1 and CTIP2 (BCL11B) (Figure 3B). However, the KCNT1 signal was negative in glial fibrillary acidic protein (GFAP) cells throughout the deep to superficial cortex (Figure 3B). These results indicated that KCNT1 is primarily expressed within deep-layer postmitotic neurons in the cortical plate, and does not appear to be expressed in progenitor or glial cells or immature neurons in any regions of the neocortical column.

**Fig. 3.**
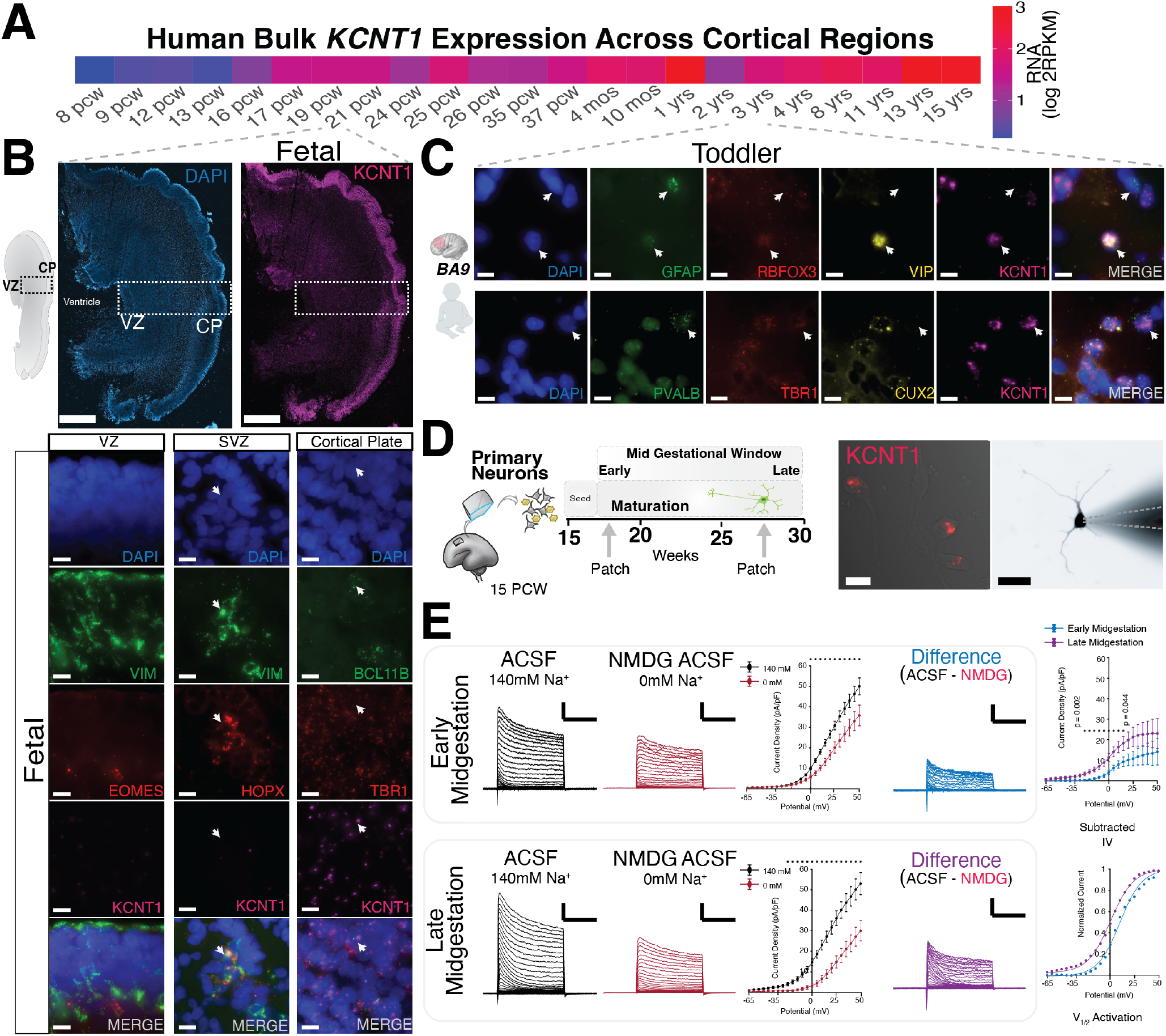
A. Analysis of KCNT1 expression during human brain development from 8 PCW to late adolescence. B. Top, KCNT1 RNA in situ hybridization of a 21 PCW human coronal brain section from the perisylvian region demonstrates robust KCNT1 enrichment in the cortical plate (CP). Bottom, spatial transcriptomic analysis of KCNT1 expression within specific CP neuronal markers, including a neuronal marker (RBFOX3) and deep layer markers CTIP2 (BCL11B) and TBR1, with limited KCNT1 expression in dividing cell types, including oRGCs and progenitor markers (HOPX, EOMES, vimentin, VIM). SP, subplate; SVZ subventricular zone; VZ, ventricular zone (scale bars: Top, 100 μm; Bottom, 10 μm). C. Analysis of KCNT1 and cell-type markers in the toddler cortex (3 years old), including enrichment in inhibitory neurons (parvalbumin and VIP expressing, PVARB, VIP) and KCNT1 expression in excitatory neurons (TBR1+ and CUX2+), with limited expression within the glia marker GFAP (scale bars: 10 μm). D. Left, experimental schematic of primary neuron plating (15 PCW) and maturation in cell culture from a healthy individual. Right, representative epifluorescence images of a primary neuron demonstrating KCNT1 expression (red signal, in situ) and a dye-loaded neuron during patch recordings (Alexa-Fluor 488 dye, 1 μM) for morphological quantification (scale bars: 10 μm). E. Representative voltage clamp recordings of outward K+ currents from mid-gestational primary human neurons evoked by an activation routine from -80 mV to +50 mV at 5 mV increments (scale bars: 1 nA/100 ms) in external ACSF with physiological sodium (140 mM) and no sodium ACSF (NMDG-ACSF, 0 mM Na+). IV plot of K+ currents in ACSF (140 mm Na+) and in NMDG-ACSF (0 mM Na+) show that KNa current partially underlies the observed increase in steady-state outward current during maturation from early to late mid-gestation. Boltzmann curves fit to the Na+-dependent outward current (total K+ current in ACSF minus the sodium-insensitive K+ current NMDG-ACSF) show a leftward shift in V1/2 activation during the mid-gestational period (95% confidence intervals for the nonlinear least-squares parameter estimates beta). Data analyzed from 18. See Table S3 for statistics.

### Increase in sodium-activated K^+^ channels (KNa1.1) in the mid-gestation human neocortex

To determine the prenatal emergence of functional KNa1.1, we performed patch-clamp recording on primary human neurons isolated from the mid-gestational neocortex (Figure 3D). We applied voltage-step protocols to elicit steady-state outward K+ channel currents over a range of membrane potentials (−80 to +60 mV). To isolate the sodium-activated K+ conductance (KNa) from the total outward steady-state K+ current, we performed patch-clamp experiments in both regular ACSF and in sodium-free ACSF in which the sodium was replaced with an equimolar concentration of N-methyl-D-glucamine (NMDG-ACSF, see Methods). To quantify the sodiumactivated K+ current, we subtracted the sodium-insensitive current (in NMDG-ACSF) from the total K+ current (in regular ACSF), leaving only the sodium-sensitive K+ current as the difference. In these experiments, the internal recording solution did not contain sodium, ensuring that KNa activation originated from sodium influx into neurons. The sodium-sensitive outward K+ current was present as early as 16 PCW (early mid-gestation) and increased in amplitude as primary neurons matured in culture over 2-3 months (Figure 3D; early vs. late, voltage steps from -25 to +15 mV, p < 0.05, n = 13 and n = 21, respectively).

KNa1.1 channels are voltage-sensitive, and epilepsy-associated variants can result in shifts in the voltage dependence of channel opening20. Therefore, we determined the half-activation voltage (V1/2) of KNa1.1 in mid-gestational primary human neurons. Performing voltage steps in both NMDG-ACSF (0 mM sodium) and regular ACSF (140 mM sodium), we isolated the V1/2 of the sodium-sensitive portion of the outward current and observed a 10-mV leftward shift in activation kinetics at later midgestational timepoints (17 PCW: V1/2 = 10.03 ± 1.70 mV; 28 PCW: V1/2 = 0.39 ± 1.14 mV, shown as 95% confidence intervals; Figure 3E). These results suggest that, in more mature neurons, the open probability of KNa1.1 is higher at more hyperpolarized potentials, resulting in KNa1.1 activation closer to Vm. Taken together, the observed increase in sodium-sensitive current and leftward shift in V1/2 activation during gestational maturation suggests that KCNT1 has a temporally precise function during development. Therefore, pathogenic variants that affect KNa1.1 biophysics might alter prenatal neurophysiology at distinct stages of development.

### KCNT1 is enriched to both inhibitory and excitatory neurons in the early childhood neocortex

Given the severity of KCNT1-associated epilepsy in infants, including 50% loss of life by 3 years of age and severe intellectual disability, we analyzed the postnatal human neocortex for KCNT1-enriched cell types. We performed multiplexed spatial RNA profiling in an infant neocortex (age: 7 months) and a toddler neocortex (age: 3 years) (Brodmann area: BA9). In the infant neocortex, KCNT1 was enriched for several neuronal subtypes, including high expression in several interneuron subtypes, such as somatostatin (SST+), vasoactive intestinal peptide (VIP+), and parvalbumin (PVALB+) subtypes (Figure S3B). Similarly, in the toddler neocortex, KCNT1 was also enriched in neurons, including the same interneuron types observed in the infant neocortex (PVALB+, SST+, VIP+; Figure 3C). Neither the infant nor toddler sample had significant KCNT1 expression in GFAP+ cells. These results suggest that a broad KCNT1 enrichment to several neuron subtypes, both excitatory and inhibitory types, during the early postnatal period likely underlies the postnatal developmental pathology.

### ASO can knock down KNa1.1 in mid-gestational human fetal neurons

Patient-derived iPSCs use NGN2 induction to generate a cortical-like excitatory neuron, yet do not fully recapitulate primary human neuron physiology. Therefore, we next investigated the feasibility of ASO knockdown of KCNT1 in primary human neurons from the midgestation neocortex. We isolated primary neurons from 15- and 23-post conception weeks neocortices and maintained the neurons in culture for 10 weeks (Figures 4, S4). We applied ASO treatment for 14 days prior to performing patch clamp recording to ensure KCNT1 knockdown. Immunohistochemical analysis using cell-type markers for neurons (MAP2 and TUJ1) and astrocytes (GFAP) demonstrated no observable changes between Vehicle neurons and neurons treated with ASO (Figure 4A). To explore the complexity and health of primary neurons following ASO treatment, we performed morphological reconstructions of patch-clamp-recorded cells and observed comparable levels of complexity (Sholl analysis, soma size, ramification index) between vehicle-, NT-ASO-, and ASO-treated cells (Figure 4B, Table S2; p > 0.873). However, NT-ASO and ASO treatment both resulted in a mild reduction of the longest neurite length (Figure 4B; Vehicle vs. ASO: p = 0.027; Vehicle vs. NT-ASO: p = 0.05), a trend also observed in the NGN2 neurons. Using voltage clamp, we demonstrated a significant ASO knock-down of steady-state outward K+ currents at depolarized potentials (25 to 60mV) compared with vehicle-and NT-ASO-treated neurons (Figure 4C; e.g., at 60 mV, vehicle: 58.02 ± 5.93 pA/pF; ASO: 43.28 ± 2.72 pA/pF; vehicle vs. ASO: p = 0.0321). Capacitance values were comparable across all three conditions (vehicle: 31.97 ± 3.20 pF; ASO: 32.63 ± 3.15 pF; NT-ASO: 27.89 ± 3.89 pF; p > 0.05). This whole-cell K+ current knockdown was observed in both fetal samples treated (Figure S4C). Taken together, these results suggest the ASO treatment can be used to knock down KNa1.1 in primary human fetal neurons.

**Fig. 4.**
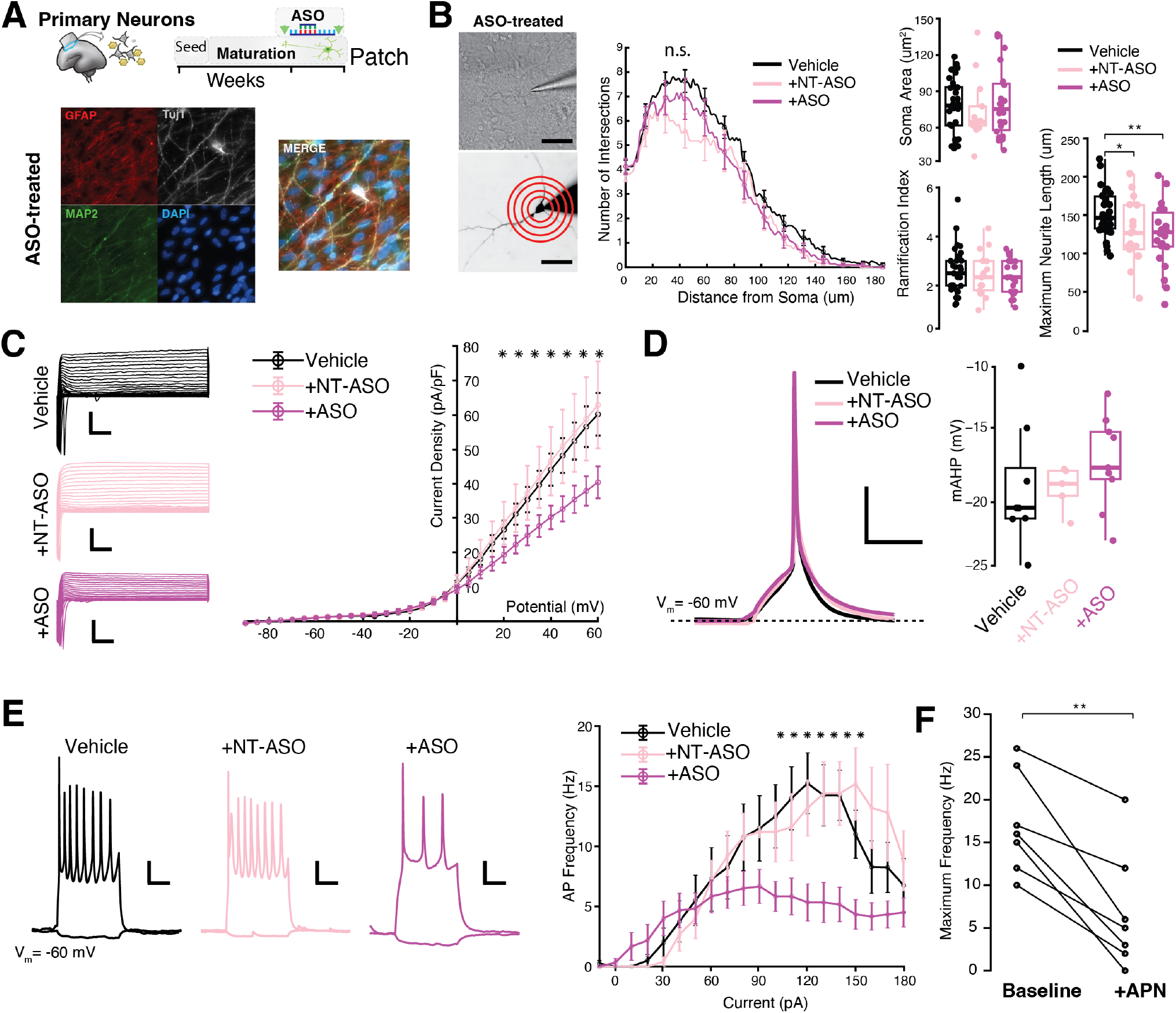
A. Top, schematic of primary neuron isolation from midgestation human cortex and maturation timeline. Bottom, Representative immunofluorescence images of primary neurons treated with ASO (14 days) and vehicle with antibody labeling of neuronal-specific markers, microtubule-associated protein 2 (MAP2) and beta-tubulin III (TUJ1), and the astrocyte marker GFAP (scale bars: 25 μm). B. Left, representative brightfield and epifluorescence images of ASO-treated primary neurons loaded with Alexa 488 dye (1 μM) with representative sholl analysis (scale bars: 50 μm). Right, analysis of neuromorphological properties of mid-gestation neurons demonstrate comparable neuronal complexity, ramification, and soma size following vehicle, NT-ASO, or ASO treatment. Maximum neurite length was significantly lower in cells treated with ASO or NT-ASO (p = 0.0089, p = 0.05, respectively). C. Representative voltage clamp traces from mid-gestation neurons evoked by step activation routine from -90 mV to +60 mV at 5 mV increments (scale bars: 1 nA/100 ms). Healthy primary fetal neurons (15 PCW sample) have steady-state outward K+ currents sensitive to ASO knockdown (10 μM for 14 days) (p = 0.0321 at 60 mV) and are unaffected by control NT-ASO (p = 0.7827 at 60 mV). D. Analysis of average first AP elicited during rheobase routine following 14-day ASO treatment. AP kinetics, including the mAHP, are unaffected by the ASO treatment, suggesting SK upregulation did not occur in primary human neurons. E. Left, representative maximum spiking of mid-gestation neurons following ASO treatment. Right, ASO-treated primary neurons displayed decreased AP firing over a range of current injections and exhibited maximum firing frequency at lower current injections than vehicle- and NT-ASO-treated cells (p = 0.012). F. Fetal neurons bath-perfused with apamin (50 nM) show reduced maximum firing frequency, suggesting SK channels help support high frequency firing in primary human neurons. * p <0.05, ** P <0.01. See Table S4 for statistics.

### ASO knockdown of KNa1.1 in mid-gestational human fetal neurons reduces excitability

To ascertain the AP firing properties of primary fetal neurons treated with ASO, we injected current (−10 to 180 pA, 10-pA steps) into neurons to generate sustained depolarizations and APs (Figure 4E). In ASO-treated primary neurons, we observed attenuated firing compared with vehicle- and NT-ASO-treated neurons. At low current inputs, ASO treatment had minimal effects (at 40 pA, vehicle: 10.55 ± 0.86 Hz, n = 10; ASO: 5.33 ± 1.85 Hz, n = 12, p = 0.023; Figure 4E); however, at large current injections, ASO-treated cells fired at lower frequencies than the vehicle- and NT-ASO-treated cells (at 80–150 pA, vehicle: 5.00 ± 0.58 Hz, n = 10; ASO: 18.22 ± 2.17 Hz, n = 12, p = 0.006; Figure 4E). Primary fetal neurons displayed significant reductions in maximum AP firing when perfused with APN (Figure 4F), suggesting that SK channels similarly support high frequency firing. Quantification of rheobase demonstrated comparable minimum necessary current for AP firing in mid-gestational primary neurons treated with Vehicle, NT-ASO, or ASO (Figure S4D, p = 0.969; see Table S4) and no significant change in AP threshold across conditions (Figure S4E, p = 0.052; see Table S4). Next, we analyzed the mAHP of ASO-treated primary neurons and found that the mAHP amplitude is comparable across treatment conditions (Figure 4D; vehicle vs. ASO: p = 0.365; see Table S4). Therefore, fetal neurons treated with ASO did not recapitulate the NGN2-neuron compensatory response of increased mAHP amplitude and increased rheobase. Taken together, our results demonstrate that ASO reduced excitability and that primary human neurons, unlike NGN2-derived neurons, respond to ASO treatment with a hypoexcitable state without any induced dependence on SK channel activation.

### Quinidine decreases the steady-state outward K+ current in primary human mid-gestational neurons

Quinidine is a nonselective ion channel antagonist that blocks potassium and sodium channels and has been shown to reduce seizures in a subset of KCNT1 patients21,22. Yet, some children do not show seizure improvements with quinidine, including the treatment-resistant KCNT1-p.R474H EIMFS individuals in our study. Nevertheless, we evaluated the effect of quinidine (QUIN) at 100 μM on primary prenatal neurons and observed a substantial decrease in the total outward steady-state K+ current following treatment (at 60 mV, baseline: 50.71 ± 6.58 pA/pF; QUIN: 14.41 ± 1.67 pA/pF; Figure S4B). These results show that quinidine reduces outward K+ conductance in mid-gestation human neurons, albeit via a non-selective manner.

## Discussion

In this study, we identified a developmental neurophysiological basis of a first-in-human RNA therapy for children with treatment-resistant KCNT1-related epilepsy. We determined that the developmental emergence of functional KNa1.1 occurs during the human mid-gestational period and demonstrated that ASOs can modulate KNa1.1 currents in primary human fetal cortical neurons. Based on our observation that the ASO can recover excitability in KCNT1-associated EIMFS patient-derived neurons, we suggest that early perinatal targeting of KNa1.1 may provide a therapeutic benefit.

Prenatal ion channel dysfunction is associated with severe neurodevelopmental disorders12, and the enrichment of KCNT1 in mid-gestation cortical plate neurons suggest that variants in this gene may result in a similar altered developmental trajectory for cortical circuits underlying migrating focal seizures seen within days of birth (as observed for one of the p.R474H patients) as well as KCNT1 postnatal microcephaly phenotypes7. To this end, we leveraged primary human neocortices to establish when functional KNa1.1 emerge. These findings can inform dosing and treatment timelines for ASO targeting. Here, mid-gestation human neurons demonstrated KNa current as early as 16 PCW, with KNa currents increasing in size and with the voltage dependence of activation (V1/2) shifting to more negative potentials as neurons mature (i.e., with a leftward shift and the KNa1.1 activating at potentials closer to resting Vm). Potential mechanisms underlying this hyperpolarizing shift could include expression of alternative KCNT1 splice isoforms, as well as differential expression of KCNT2, which forms heteromers with KCNT123,24. However, unlike other ion channels enriched in prenatal cells (e.g., SCN3A, GRIN2B), KCNT1 was not found in neocortical progenitors, key cell types associated with brain malformations (e.g., outer radial glial cells)12, which suggests the prenatal deficits of KCNT1 are primarily neuronal.

KNa1.1 dysfunction is associated with treatment-resistant epilepsy immediately following birth in the neonatal period (e.g., EIMFS), with a high degree of heterogeneity in seizure types and inconsistent seizure reduction by various drugs25. For example, quinidine is a nonselective blocker that can reduce seizure burden in some individuals with KCNT1 GoF variants, yet fails to normalize developmental recovery21,26 and has adverse cardiac effects22. Therefore, even with early genetic identification of EIMFS, neonatal epilepsies may still have poor outcomes with the use of available drugs15, necessitating a shift from nonselective to targeted therapies27. Our results suggest that ASO targeting of perinatal KNa1.1 GoF current might help limit disease progression compared with postnatal treatment. Taking this finding together with our observed fetal cell-type enrichments, we believe that in utero delivery of RNA therapeutics may be a viable treatment modality, similar to prenatal ASO treatment of Angelman syndrome mice that improves phenotypes28, and other prenatal treatments for non-brain human diseases29.

In KCNT1 EIMFS neurons, GoF KNa results in a blunted excitability curve (neurons cannot maintain APs at high current injections), which is improved with ASO treatment, likely by increased AHP amplitude that would otherwise be inhibited by the prolonged current injection during the burst. Rodent electrophysiology data have shown a similar paradoxical finding, with GoF KCNT1 variants displaying lower excitability in cortical GABAergic neurons, resulting in a disrupted excitatory/inhibitory balance, and generating network hyperexcitability30. ASO-treated EIMFS neurons also display improved AP amplitude run-down between spikes and reduced depolarization block, demonstrating a recovery of AP kinetics during AP bursts. Several functions of KNa activation in neurons have been proposed, including shaping of complex and simple spiking to shift cellular excitability and inter-burst timing10,31,32 and enlarging AHP after a single spike to enhance burst firing during prolonged current stimulation13,33. Despite these potential actions of KNa affecting EIMFS neuronal excitability and network excitability in vitro and in vivo, we found that ASO treatment of KCNT1 and control NGN2 neurons showed an unexpected compensation: KCNT1 knockdown increased AHP amplitude by potentiation of Ca++-activated K+ channel (BK and SK) activity. The possible upregulation of SK channels in ASO-treated neurons may assist in preventing depolarization block, allowing firing at high current injections. Moreover, when APN was perfused in ASO-treated neurons, SK channels could not support high frequency firing, with cells reaching depolarization block sooner. Although the mechanism for ASO upregulation of Ca++-activated K+ channels is unclear, it is well documented that ion channel expression increases (or decreases) as a compensatory change in many ion channel families34,35. An additional potential homeostatic mechanism is the activity-dependent alternative splicing of BK channels to regulate cell and network activity36. Overall, ASO enhanced the excitability of EIMFS neurons to sustain a higher frequency of spiking activity at higher input currents. These findings highlight the ability of AHP modulation to shape cellular physiology, and future studies exploring therapeutic AHP enhancement, particularly in vivo to analyze complex spiking activity, might provide a basis for understanding a broad range of disorders with disrupted cellular excitability.

While human iPSC-induced neurons (i.e., NGN2 induction) are valuable for testing and validating targeted RNA therapeutic strategies, NGN2 overexpression creates an artificial expression system with a transcriptome and neurophysiology that does not fully match any single cell type in the human brain, yet transcriptionally most closely resembles cortical excitatory neurons37. For example, in this context, compared with the primary human neurons utilized in this study, NGN2 neurons have larger KNa1.1 and do not possess the correct protein co-expression network. Therefore, studies of early-onset disorders should ideally be validated in primary human tissue. In this work, we found that ASO treatment of primary human neurons reduced excitability at high current injections and did not modulate mAHP, in contrast to NGN2 neurons (both patient and control cell lines). Follow-up studies should also consider non-conducting roles of KCNT1 in NGN2 and primary human fetal neurons, including RNA translation, cellular signaling, and modulation by voltage-gated sodium channels38, as well as potential differential pathogenic KNa1.1 expression in the mitochondrial membrane39. Exploring downstream or alternative disease mechanisms could provide additional insights into the role of KCNT1 in normal development and could inform the design of biomarkers, enabling the recovery of developmental delay without requiring the targeting of KNa1.1 GoF currents.

The KCNT1 enrichment observed in inhibitory neurons (most abundant in PVALB+) from the infant and toddler human brain likely supports a key physiological role for KNa1.1 in GABAergic refinement of neocortical circuits, a proposed mechanism responsible for EIMFS8. Mouse models of infantile and epileptic spasms syndrome show altered Pvalb+ interneuron development and GABAergic synaptic dysfunction throughout life40, with Pvalb+ interneurons failing to fire repetitively at large amplitude current injections and prone to depolarization blockage in KCNT1-L456F mice41. Moreover, differential pathophysiology exists even within related GABAergic subtypes (SST, PVALB, VIP), resulting in opposing functional deficits42. In this context, a previous study showed that ASO downregulation in KCNT1 p.P924L GoF mice is safe and effective in controlling seizures, with improvements in cognition and behavior in presymptomatic neonatal mice43. In the context of our work with human neurons, a shared mechanism may facilitate enhanced network inhibition and input–output function by KNa1.1 enrichment in inhibitory neurons.

While we strive for a clinical path forward for KCNT1-targeted treatments, our studies indicate that KCNT1 may contribute to prenatal human brain development. Additionally, we have described the electrophysiological mechanisms by which KCNT1-knockdown ASOs can affect AP generation and neuronal excitability. These findings warrant future studies to explore the use of ASO in early perinatal clinical treatments for KCNT1-associated disorders to provide early neuroprotection against KNa1.1-GoF-associated encephalopathy and developmental delay.

## Methods

### Human subjects and samples

Human subject research was conducted according to protocols approved by the institutional review boards of Northwestern University and Boston Children’s Hospital. Fetal brain tissue was received after release from clinical pathology, with a maximum postmortem interval of 4 h. Fetal cases with known anomalies were excluded. Tissue was transported in ice-cold Hibernate-E medium (Thermo Fisher) for processing in the laboratory. The neonatal (7 months) and 3-year-old brain samples were obtained from the University of Maryland Brain and Tissue Bank of the NIH NeuroBioBank (sample numbers UMBN 4353 and HCT17HEIA029, respectively) and stored at -80°C until processing.

### Human-derived induced pluripotent stem cell lines with p.R474H-KCNT1

Patient iPSCs were derived from skin fibroblasts and reprogrammed via Sendai virus with a normal karyotype, as previously described4. iPSC colonies were maintained in mTESR1 plus media (Stem Cell Technologies) on Geltrex (Thermo Fisher)-coated plates and passaged every 5–7 days, and the media was changed every other day. The iPSCs were tested as mycoplasma-negative. Pluripotency was confirmed with qRT-PCR, and markers OCT4, NANOG, and TRA1-60 were detected using immunocytochemistry. Publicly available iPSC lines reprogrammed with Sendai virus were used for wild-type comparisons, including Personal Genome Project cell line 1 (PGP1) and the NIH reference/control cell line KOLF2.1J44. Although both KOLF2.1J and PGP1 control lines were used, after discovering potential CNV issues with KOLF2.1J, PGP1 was primarily used for studies.

### Neurogenin 2 differentiation of iPSCs

Cortical neurons were differentiated via lentiviral transduction of a tetracycline-inducible NGN2 cassette, followed by doxycycline induction (6 days) and neural induction by adding SMAD inhibitors SB431542 and XAV939 (Tocris). The medium was supplemented with BDNF (Millipore), NT3 (Peprotech), and laminin37. All experiments were repeated at least two times with independent differentiation batches of neurons, and the number of replicates is detailed after each experiment.

### Primary neuronal cultures and neurite analysis

Isolation of primary human neuron cultures was modified from a previous method45. Briefly, immediately following tissue release from pathology, the neocortex was dissected from two human samples (15 weeks and 23 weeks post-conception) according to the manufacturer’s protocol (MACs Neural Dissociation Kit, Miltenyi Biotec). Cortical neurons were plated onto laminin-treated coverslips (10 μg/ml) at 250k cells/well in 24-well plates and maintained in Neurobasal Plus media, B27 Plus, and 1:100 pen/strep (Thermo Fisher) supplemented with 1:5,000 laminin at each media change. Half-media changes were performed every 2–3 days. Neurite lengths from the soma were traced and measured using ImageJ software. Neurons were fixed with 4% PFA in PBS for 8 minutes at 4 degrees Celsius. Cells were blocked in 10% donkey serum in PBS with 0.3% Triton X-100 for 1 hour at room temperature, followed by immunostaining.

### Antisense oligonucleotide design, production, and treatment

Gapmer ASOs designed to be complementary to the KCNT1 gene (ASO), or control NT-ASO with comparable chemistry, were synthesized by Microsynth. KCNT1-specific ASO has a 5-10-5 gapmer design with a mixed phosphodiester/phosphorothioate backbone, GToToGoCCTTTGTAGCTGoAoGoGT, without any off-targets in the human transcriptome. The NT-ASO shared a similar 5-10-5 gapmer design, also without any targets in the human transcriptome. As previously described, for NGN2 neurons, ASOs were delivered via gymnosis at 10 μM (free uptake after direct addition of ASOs to culture media) from DIV 14 to DIV 21, followed by 5 μM ASO treatment from DIV 21 to DIV 28, when patching experiments began. As shown, this treatment significantly reduced KCNT1 mRNA levels. ASO incubation was maintained via half-media change for 10–14 days. In the case of primary human neurons, from DIV 21 to DIV 28, 10 *μ*M was maintained instead of 5 *μ*M to ensure robust KCNT1 knockdown.

### RNA isolation, cDNA synthesis, qRT-PCR

RNA isolation from iPSC-derived NGN2 neurons was performed using the RNeasy Plus Mini Kit (Qiagen) following the manufacturer’s protocol. The concentration and purity of the RNA samples were quantified using the Agilent Synergy LX Take 3 plate. cDNA synthesis was performed using SuperScript IV (Invitrogen) with 100 ng of RNA per reaction. The qRT-PCR protocol was performed with SYBR Green 2x Master Mix (Applied Biosystems), with forward and reverse primers designed for the target gene sequences in purified water (Ambion) on MicroAmp Endura Plate Optical 96-Well plates (Applied Biosystems). The efficiency of the primers was verified with melting curve analyses, and the qPCR reaction was conducted with the QuantStudio 6 Pro (Applied Biosystems). Quantification of KCNT1 mRNA expression was performed by normalizing to the endogenous control gene and vehicle-treated cells (mock transfection with water) using the delta-delta Ct method. Delta Ct = CtKCNT1 - CtB-actin; delta-delta Ct = CtASO - Ctvehicle; fold change = 2(−delta-delta Ct). The fold change, expressed as a percentage, served as a quantitative measurement of KCNT1 mRNA expression.

### Fluorescent immunohistochemistry

NGN2 and primary human neurons were analyzed with immunocytchemical staining to confirm cell types and distributions. For immunocytochemical characterization, the cells were first fixed with 4% PFA for 8–20 min at 4 degrees Celsius. After washing with PBS, non-specific staining was blocked with 10% BSA-PBS - 0.3% TritonX-100, and primary antibodies were diluted into 10% BSA-PBS-0.1% TritonX-100 and incubated overnight at +4°C. See the key resource table of primary antibodies used and their concentrations. After PBS washes, cells were incubated with secondary antibodies (1:500) for 1 hour at room temperature and washed three times. On the third wash, nuclei were stained with DAPI (Thermo Fisher Scientific) for 5 min at room temperature.

### Patch-clamp of iPSC-derived NGN2 and primary human neurons

Patch-clamp recordings were performed on DIV 26-32. NGN2 neurons were visualized using an upright Olympus BX51 microscope with epifluorescence. Current- and voltage-clamp data were recorded and analyzed with SutterPatch IPA (Sutter Instruments, Sunnyvale, CA). Glass pipettes were formed from borosilicate glass (3–5 MO) on a Sutter Puller and filled with a no-sodium intracellular solution to isolate sodium-activated currents; 97.5 mM K-Gluc, 32.5 mM KCl, 10 mM HEPES, 1 mM EGTA, and 2 mM MgCl2. The internal solution was adjusted to 295 mOsm and a pH of 7.4 with KOH. The internal solution also contained Alexa-Fluor 488 nm dye to visualize neuron morphology (1 mM, Thermo Fisher). For patching of NGN2 neurons, the extracellular recording solution, artificial cerebral spinal fluid (ACSF), contained the following: 125mM NaCl, 2.5mM KCl, 2mM CaCl2, 1mM MgCl2, 1.25mM NaH2PO4, 26mM NaHCO3, 15mM glucose, 1 mM Myo-inositol, 2 mM Na-pyruvate, 0.4 mM ascorbic acid, pH to 7.4 with NaOH, 300mOsm. ACSF solution was continuously oxygenated during experiments with 95% O2 and 5% CO2 to maintain pH 7.4. For patching of fetal neurons, the extracellular recording solution, ACSF, contained the following: 140 mM NaCl, 5.4 mM KCl, 1 mM CaCl2, 1 mM MgCl2, 10 mM HEPES, and 10 mM Glucose, pH to 7.4 with NaOH, 300 mOsm. The sodium substitution external solution (NMDG-ACSF) contained 140 mM NMDG, 5.4 mM KCl, 1 mM CaCl, 1 mM MgCl2, 10 mM HEPES, and 10 mM Glucose. For the number of cells patched per genotype, see Tables S1–S4.

For voltage-clamp recordings on iPSC-derived neurons, 500 ms voltage pulses beginning at -80 mV and ending at +50 mV (10 mV increments) were used to elicit currents. The outward steady-state current was measured in response to each voltage pulse as the mean of the last 10 ms of the response. For primary cortical neurons, the voltage-clamp routine was slightly modified to begin at -90 mV (instead of -80 mV) and progress in 5 mV increments up to +60 mV to better determine the half-activation potential of KCNT1.

To study the first action potential kinetics in current-clamp (rheobase), 40 ms current steps increasing in 2 pA increments were injected into the neuron until the first action potential was achieved. The first action potential threshold was calculated by locating the point during depolarization where the first derivative of the voltage vs. time plot was minimized within a 15 ms window before the peak. Action potential amplitude was calculated as the potential difference between the threshold and the peak of the AP. The mAHP was defined as the potential difference between the threshold and the voltage minimum within a 200 ms window after the peak of the AP. The full width of the action potential was defined as the horizontal (temporal) distance between the upstroke and downstroke of the AP at the threshold potential. AP latency was defined as the time interval between the beginning of the current pulse and the time at which the AP threshold was reached. The mAHP area was calculated using a trapezoidal Riemann sum with a step size of Δ*t* = 0.1 ms with respect to the holding potential (V_m_ = -60 mV).

### Human brain tissue preparation and spatial mRNA in situ hybridization

Brain tissue preparation and spatial mRNA in situ hybridization were performed as previously described by Smith et al19,45. Briefly, following fixation (4% PFA) and cryoprotection (30% sucrose), brains were frozen using isopentane on dry ice. Samples were sectioned at 8–20 μm thickness using a Leica Cryostat, mounted immediately onto warm charged SuperFrost Plus slides (Fisher), and stored at –80°C. Fluorescent imaging was performed on selected cortical areas using an Axio Imager M2 microscope (Zeiss, Germany) or a Nikon S10 microscope. The cells were imaged with a 40X objective. We followed the manufacturer’s standard protocol for multiplex fluorescent in situ hybridization (Multiplex Version 2 kit, Advanced Cell Diagnostics). See the supplemental materials for in situ probe ACD catalog numbers. Bright-field and fluorescent images were background-corrected using Zen Blue Software for center intensity illumination and stitched together. Analysis was performed by quantifying KCNT1 puncta in cell types defined by the presence of at least 10 puncta of specific markers (Figure S3).

### Bulk human cortex gene expression analysis

The Allen Brain Atlas publishes a rich dataset of cortical gene expression across cortical brain regions from age 8 weeks post-conception to adulthood18. BrainSpan data analysis of KCNT1 (chr1:116,915,289-116,952,883, GRCh37/hg19) was performed. RNA-seq expression, measured in RPKM (reads per kilobase exon per million mapped reads), was obtained from the BrainSpan project data and summarized to Gencode v10 exons for all annotated neocortical tissues aged 12 weeks post-conception to 36 years. Brain regions for Figure 3 include the dorsolateral prefrontal cortex, ventrolateral prefrontal cortex, anterior (rostral) cingulate (medial prefrontal) cortex, orbital frontal cortex, primary motor-sensory cortex, parietal neocortex, posterior (caudal) superior temporal cortex (area 22c), inferolateral temporal cortex (area 20), occipital neocortex, temporal neocortex, primary motor cortex (area 4), primary somatosensory cortex (area S1, areas 3, 1, 2), posteroventral (inferior) parietal cortex, and primary auditory cortex (core).

### Statistical analyses

Statistical analyses were performed using MATLAB and/or R. Depending on normality, parametric or non-parametric tests were conducted: two-way ANOVA followed by Tukey’s multiple comparisons test or Kruskal–Wallis test followed by Dunn’s multiple comparisons test, with p-values adjusted for multiple comparisons. For independent or within-group comparisons, unpaired ttests (parametric and non-parametric) with one-tailed and two-tailed p-values were used. Unless otherwise indicated, p-values are presented as follows: *p<0.05, **p<0.01, ***p<0.001, and ****p<0.0001. Figure legends include the details of the statistical methods used for the analysis of each dataset. Boxplots show the median values +/-interquartile range. Table values are mean +/-SEM and p-values are reported between Vehicle and ASO-treated condition unless otherwise noted.

## Supporting information

Supplemental Results

## Author Contributions

R.S.S., S.R.G., and K.S. contributed to experimental planning, neuronal differentiation experiments, and the writing of the manuscript. R.S.S. and S.R.G. performed patch clamp recordings, pharmacology, and data analyses. R.H. provided guidance on electrophysiology experimental planning and analysis. K.S. and M.C.B. performed the neuronal differentiation experiments, qRT-PCR expression analyses, and tissue-based work. K.S. and A.C.B. assisted with imaging analysis, RNA stainings, and statistics. T.W.Y. and T.N. contributed the iPSC lines and ASOs. All authors read and edited the manuscript.

## Acknowledgements

We are grateful to the families for participating in this research. We thank Dina Simkin and Al George for project feedback. This work was supported by R00NS112604 (R.S.S.).

---

**Table S1:**
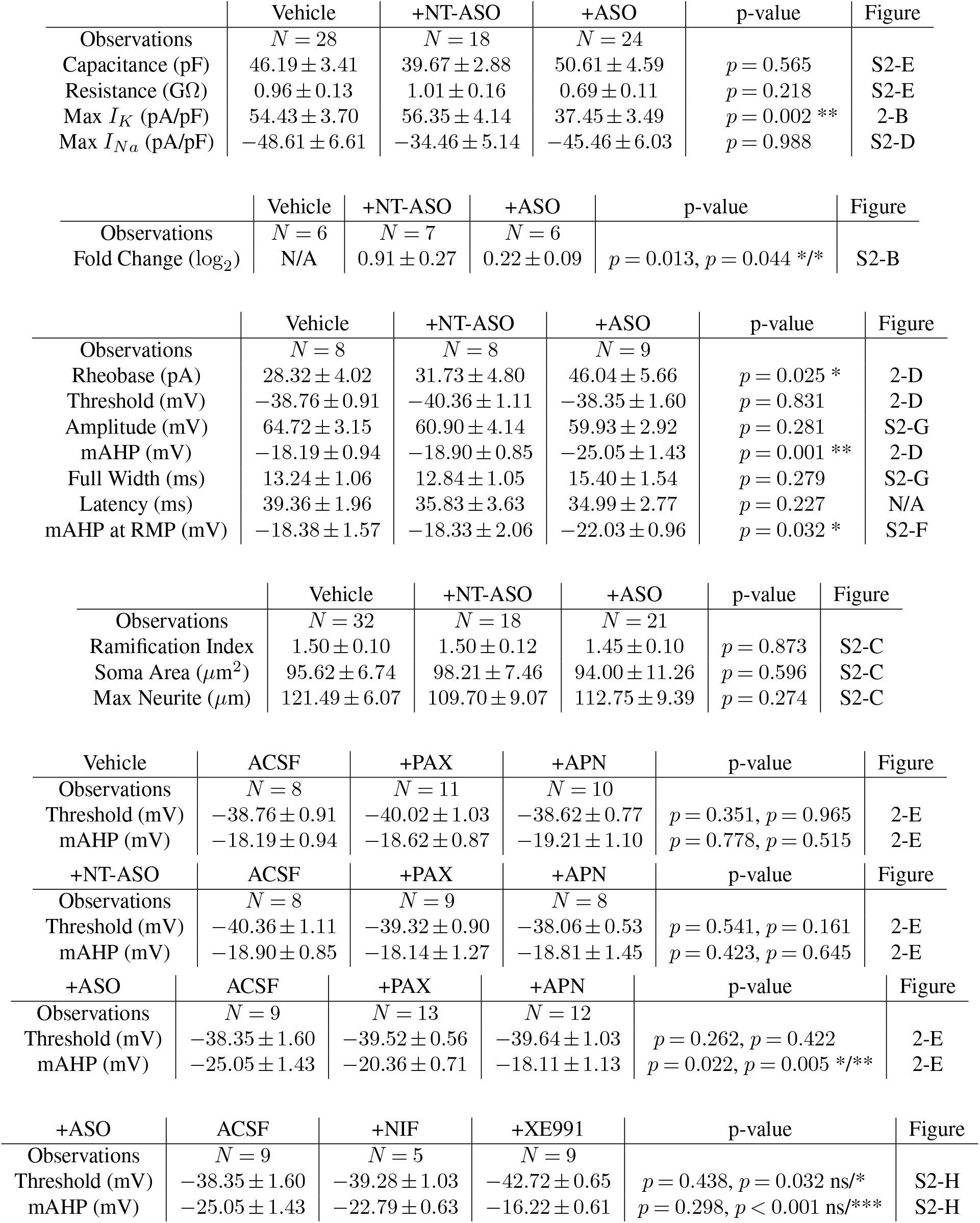
Summary of voltage clamp and action potential kinetics of KCNT1-p.R474H neurons.

| +ASO | ACSF | +NIF | +XE991 | p-value | Figure |
| --- | --- | --- | --- | --- | --- |
| Observations | $N = 9$ | $N = 5$ | $N = 9$ | | |
| Threshold (mV) | $-38.35 \pm 1.60$ | $-39.28 \pm 1.03$ | $-42.72 \pm 0.65$ | $p = 0.438, p = 0.032 \text{ ns}/^{*}$ | S2-H |
| mAHP (mV) | $-25.05 \pm 1.43$ | $-22.79 \pm 0.63$ | $-16.22 \pm 0.61$ | $p = 0.298, p < 0.001 \text{ ns}/^{***}$ | S2-H |

**Table S2:**
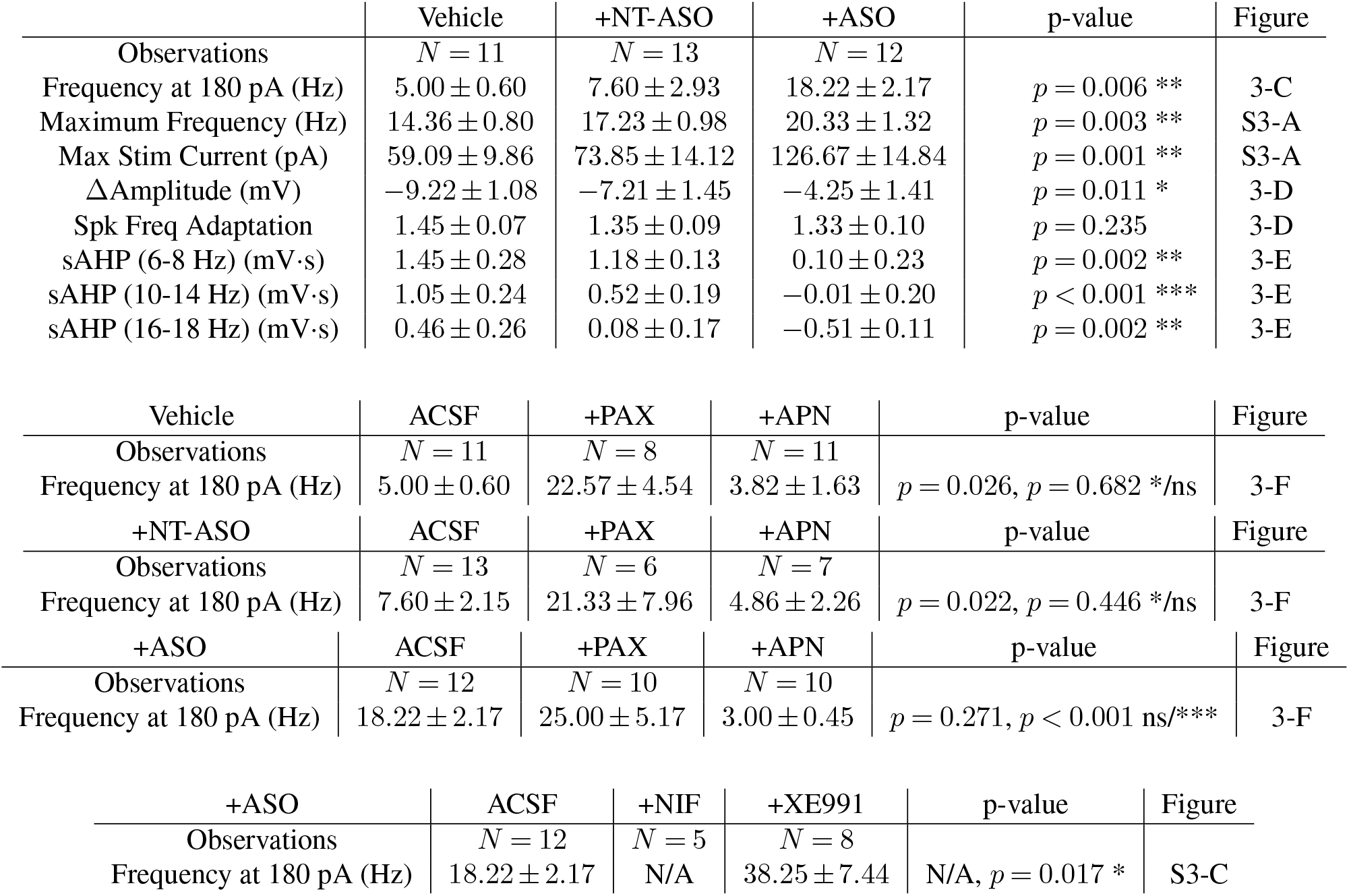
Summary of burst firing and sAHP of KCNT1-p.R474H neurons.

|  | Vehicle | +NT-ASO | +ASO | p-value | Figure |
| --- | --- | --- | --- | --- | --- |
| Observations | *N* = 11 | *N* = 13 | *N* = 12 |  |  |
| Frequency at 180 pA (Hz) | 5.00 ± 0.60 | 7.60 ± 2.93 | 18.22 ± 2.17 | *p* = 0.006 ** | 3-C |
| Maximum Frequency (Hz) | 14.36 ± 0.80 | 17.23 ± 0.98 | 20.33 ± 1.32 | *p* = 0.003 ** | S3-A |
| Max Stim Current (pA) | 59.09 ± 9.86 | 73.85 ± 14.12 | 126.67 ± 14.84 | *p* = 0.001 ** | S3-A |
| ΔAmplitude (mV) | −9.22 ± 1.08 | −7.21 ± 1.45 | −4.25 ± 1.41 | *p* = 0.011 * | 3-D |
| Spk Freq Adaptation | 1.45 ± 0.07 | 1.35 ± 0.09 | 1.33 ± 0.10 | *p* = 0.235 | 3-D |
| sAHP (6-8 Hz) (mV·s) | 1.45 ± 0.28 | 1.18 ± 0.13 | 0.10 ± 0.23 | *p* = 0.002 ** | 3-E |
| sAHP (10-14 Hz) (mV·s) | 1.05 ± 0.24 | 0.52 ± 0.19 | −0.01 ± 0.20 | *p* < 0.001 *** | 3-E |
| sAHP (16-18 Hz) (mV·s) | 0.46 ± 0.26 | 0.08 ± 0.17 | −0.51 ± 0.11 | *p* = 0.002 ** | 3-E |

| Vehicle | ACSF | +PAX | +APN | p-value | Figure |
| --- | --- | --- | --- | --- | --- |
| Observations | *N* = 11 | *N* = 8 | *N* = 11 |  |  |
| Frequency at 180 pA (Hz) | 5.00 ± 0.60 | 22.57 ± 4.54 | 3.82 ± 1.63 | *p* = 0.026, *p* = 0.682 */ns | 3-F |

| +NT-ASO | ACSF | +PAX | +APN | p-value | Figure |
| --- | --- | --- | --- | --- | --- |
| Observations | *N* = 13 | *N* = 6 | *N* = 7 |  |  |
| Frequency at 180 pA (Hz) | 7.60 ± 2.15 | 21.33 ± 7.96 | 4.86 ± 2.26 | *p* = 0.022, *p* = 0.446 */ns | 3-F |

| +ASO | ACSF | +PAX | +APN | p-value | Figure |
| --- | --- | --- | --- | --- | --- |
| Observations | *N* = 12 | *N* = 10 | *N* = 10 |  |  |
| Frequency at 180 pA (Hz) | 18.22 ± 2.17 | 25.00 ± 5.17 | 3.00 ± 0.45 | *p* = 0.271, *p* < 0.001 ns/*** | 3-F |

| +ASO | ACSF | +NIF | +XE991 | p-value | Figure |
| --- | --- | --- | --- | --- | --- |
| Observations | *N* = 12 | *N* = 5 | *N* = 8 |  |  |
| Frequency at 180 pA (Hz) | 18.22 ± 2.17 | N/A | 38.25 ± 7.44 | N/A, *p* = 0.017 * | S3-C |

**Table S3:**
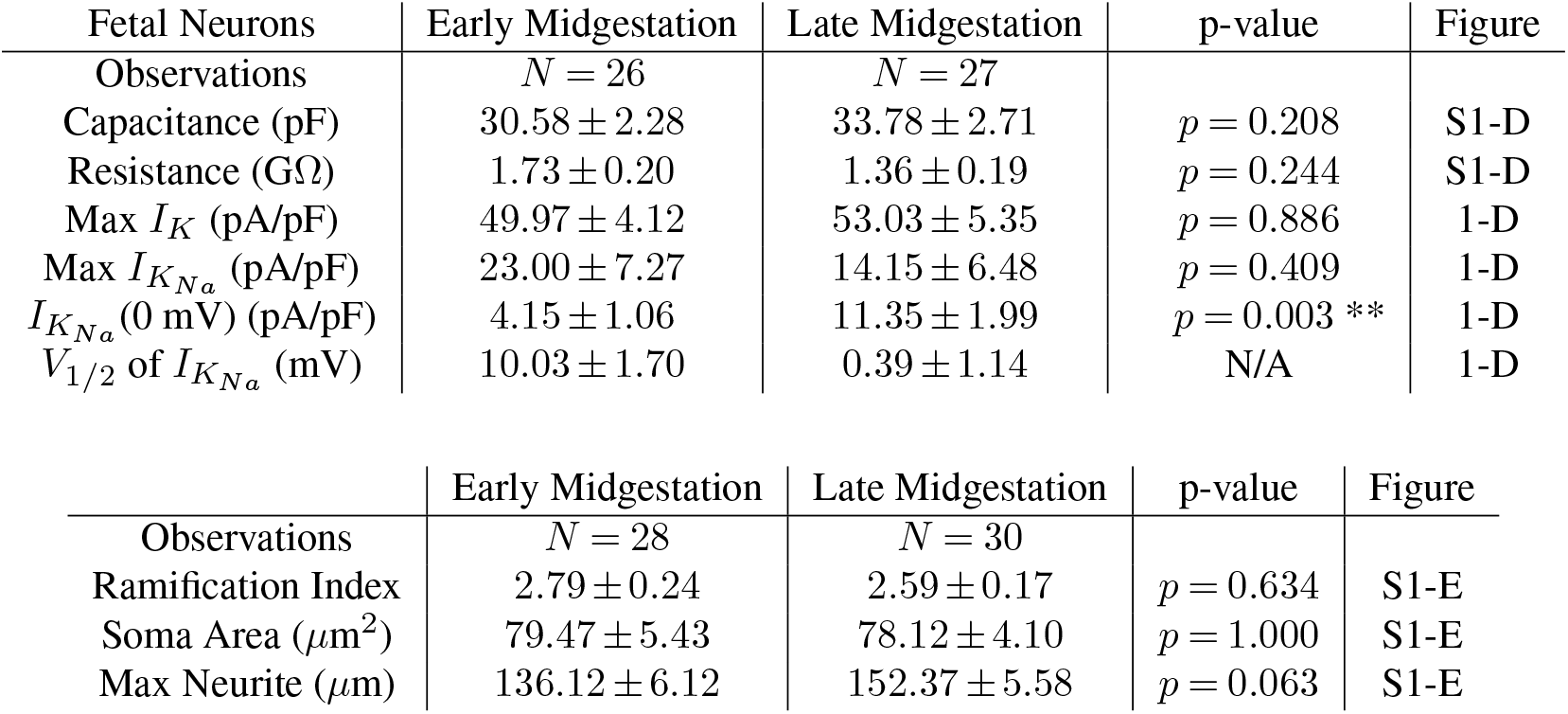
Summary of early and late mid-gestation primary fetal neurons.

| Fetal Neurons | Early Midgestation | Late Midgestation | p-value | Figure |
| --- | --- | --- | --- | --- |
| Observations | *N* = 26 | *N* = 27 |  |  |
| Capacitance (pF) | 30.58 ± 2.28 | 33.78 ± 2.71 | *p* = 0.208 | S1-D |
| Resistance (GΩ) | 1.73 ± 0.20 | 1.36 ± 0.19 | *p* = 0.244 | S1-D |
| Max *I<sub>K</sub>* (pA/pF) | 49.97 ± 4.12 | 53.03 ± 5.35 | *p* = 0.886 | 1-D |
| Max *I<sub>KNa</sub>* (pA/pF) | 23.00 ± 7.27 | 14.15 ± 6.48 | *p* = 0.409 | 1-D |
| *I<sub>KNa</sub>* (0 mV) (pA/pF) | 4.15 ± 1.06 | 11.35 ± 1.99 | *p* = 0.003 ** | 1-D |
| *V<sub>1/2</sub>* of *I<sub>KNa</sub>* (mV) | 10.03 ± 1.70 | 0.39 ± 1.14 | N/A | 1-D |

|  | Early Midgestation | Late Midgestation | p-value | Figure |
| --- | --- | --- | --- | --- |
| Observations | *N* = 28 | *N* = 30 |  |  |
| Ramification Index | 2.79 ± 0.24 | 2.59 ± 0.17 | *p* = 0.634 | S1-E |
| Soma Area (μm<sup>2</sup>) | 79.47 ± 5.43 | 78.12 ± 4.10 | *p* = 1.000 | S1-E |
| Max Neurite (μm) | 136.12 ± 6.12 | 152.37 ± 5.58 | *p* = 0.063 | S1-E |

**Table S4:**
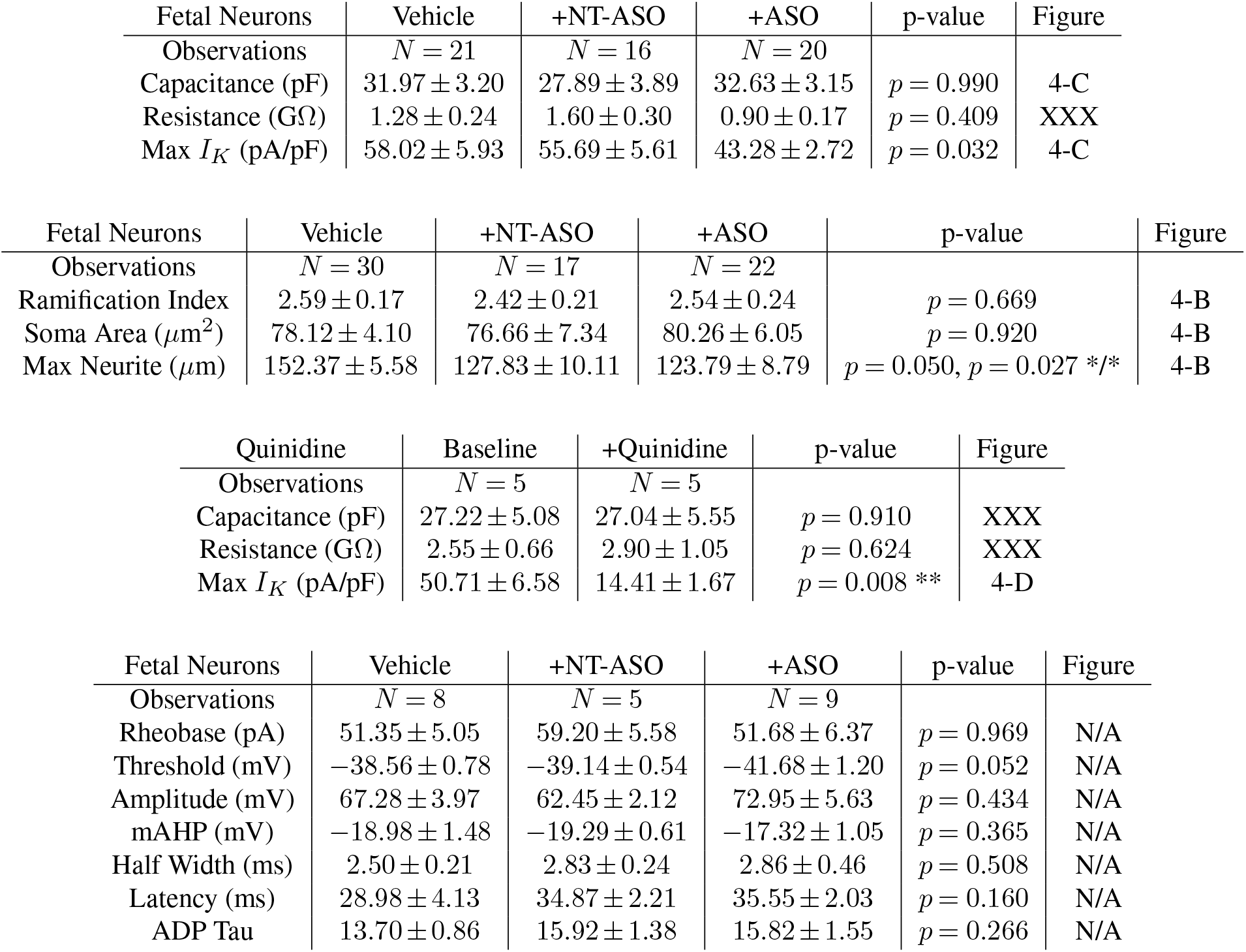
Summary of ASO treatment conditions of primary fetal neurons.

| Fetal Neurons | Vehicle | +NT-ASO | +ASO | p-value | Figure |
| --- | --- | --- | --- | --- | --- |
| Observations | $N = 8$ | $N = 5$ | $N = 9$ | | |
| Rheobase (pA) | $51.35 \pm 5.05$ | $59.20 \pm 5.58$ | $51.68 \pm 6.37$ | $p = 0.969$ | N/A |
| Threshold (mV) | $-38.56 \pm 0.78$ | $-39.14 \pm 0.54$ | $-41.68 \pm 1.20$ | $p = 0.052$ | N/A |
| Amplitude (mV) | $67.28 \pm 3.97$ | $62.45 \pm 2.12$ | $72.95 \pm 5.63$ | $p = 0.434$ | N/A |
| mAHP (mV) | $-18.98 \pm 1.48$ | $-19.29 \pm 0.61$ | $-17.32 \pm 1.05$ | $p = 0.365$ | N/A |
| Half Width (ms) | $2.50 \pm 0.21$ | $2.83 \pm 0.24$ | $2.86 \pm 0.46$ | $p = 0.508$ | N/A |
| Latency (ms) | $28.98 \pm 4.13$ | $34.87 \pm 2.21$ | $35.55 \pm 2.03$ | $p = 0.160$ | N/A |
| ADP Tau | $13.70 \pm 0.86$ | $15.92 \pm 1.38$ | $15.82 \pm 1.55$ | $p = 0.266$ | N/A |

## Bibliography

1. Finkel, R. S. et al. Nusinersen versus Sham Control in Infantile-Onset Spinal Muscular Atrophy. N. Engl. J. Med. 377, 1723–1732 (2017).

2. Korobeynikov, V. A., Lyashchenko, A. K., Blanco-Redondo, B., Jafar-Nejad, P. Shneider, N. A. Antisense oligonucleotide silencing of FUS expression as a therapeutic approach in amyotrophic lateral sclerosis. Nat. Med. 28, 104–116 (2022).

3. Tran, H. et al. Suppression of mutant C9orf72 expression by a potent mixed backbone antisense oligonucleotide. Nat. Med. 28, 117–124 (2022).

4. Nakayama et al. Antisense oligonucleotide therapy for KCNT1 epilepsy of infancy with migrating focal seizures. Under review (2024).

5. Kuchenbuch, M. et al. KCNT1 epilepsy with migrating focal seizures shows a temporal sequence with poor outcome, high mortality and SUDEP. Brain 142, 2996–3008 (2019).

6. Burgess, R. et al. The Genetic Landscape of Epilepsy of Infancy with Migrating Focal Seizures. Ann. Neurol. 86, 821–831 (2019).

7. Barcia, G. et al. De novo gain-of-function KCNT1 channel mutations cause malignant migrating partial seizures of infancy. Nature Publishing Group 44, 1255–1259 (2012).

8. Kim, G. E. et al. Human Slack Potassium Channel Mutations Increase Positive Cooperativity between Individual Channels. Cell Rep. 9, 1661–1672 (2014).

9. Kaczmarek, L. K. Slack, Slick, and Sodium-Activated Potassium Channels. ISRN Neurosci. 2013, 354262 (2013).

10. Yang, B., Desai, R. Kaczmarek L.K. Slack and Slick K Na Channels Regulate the Accuracy of Timing of Auditory Neurons. J. Neurosci. 27, 2617–2627 (2007).

11. Milligan, C. J. et al. KCNT1 gain of function in 2 epilepsy phenotypes is reversed by quinidine. Ann. Neurol. 75, 581–590 (2014).

12. Smith, R.S. Walsh, C.A. Ion Channel Functions in Early Brain Development. Trends in Neurosciences 43, 103–114 (2020).

13. Franceschetti, S. et al. Na+-Activated K+ Current Contributes to Postexcitatory Hyperpolarization in Neocortical Intrinsically Bursting Neurons. J. Neurophysiol. 89, 2101–2111 (2003).

14. Andrade, R., Foehring, R.C. Tzingounis, A.V. The calcium-activated slow AHP: cutting through the Gordian knot. Front. Cell. Neurosci. 6, 47 (2012).

15. Berg, A. T. et al. Immediate outcomes in early life epilepsy: A contemporary account. Epilepsy Behav. 97, 44–50 (2019).

16. McTague, A. et al. Clinical and molecular characterization of KCNT1-related severe early-onset epilepsy. Neurology 90, 10.1212/WNL.0000000000004762 (2017).

17. McTague, A. et al. Migrating partial seizures of infancy: expansion of the electroclinical, radiological and pathological disease spectrum. Brain 136, 1578–1591 (2013).

18. Jones, A., Overly, C.C. Sunkin, S.M. The Allen Brain Atlas: 5 years and beyond. Nat Rev Neurosci 10, 821–828 (2009).

19. Smith, R. S. et al. Early role for a Na+,K+-ATPase (ATP1A3) in brain development. Proc. Natl. Acad. Sci. U.S.A. 118, (2021).

20. Rychkov, G. Y. et al. Functional Effects of Epilepsy Associated KCNT1 Mutations Suggest Pathogenesis via Aberrant Inhibitory Neuronal Activity. Int. J. Mol. Sci. 23, 15133 (2022).

21. Dilena, R. et al. Early Treatment with Quinidine in 2 Patients with Epilepsy of Infancy with Migrating Focal Seizures (EIMFS) Due to Gain-of-Function KCNT1 Mutations: Functional Studies, Clinical Responses, and Critical Issues for Personalized Therapy. Neurotherapeutics 15, 1112–1126 (2018).

22. Mikati, M. A. et al. Quinidine in the treatment of KCNT1-positive epilepsies. Ann. Neurol. 78, 995–999 (2015).

23. Chen, H. et al. The N-Terminal Domain of Slack Determines the Formation and Trafficking of Slick/Slack Heteromeric Sodium-Activated Potassium Channels. J. Neurosci. 29, 5654–5665 (2009).

24. Joiner, W. J. et al. Formation of intermediate-conductance calcium-activated potassium channels by interaction of Slack and Slo subunits. Nat. Neurosci. 1, 462–469 (1998).

25. Borlot, F. et al. KCNT1-related epilepsy: An international multicenter cohort of 27 pediatric cases. Epilepsia 61, 679–692 (2020).

26. Numis, A. L. et al. Lack of response to quinidine in KCNT1-related neonatal epilepsy. Epilepsia 59, 1889–1898 (2018).

27. Bonardi, C. M. et al. KCNT1 -related epilepsies and epileptic encephalopathies: phenotypic and mutational spectrum. Brain 144, 3635–3650 (2021).

28. Clarke, M. T. et al. Prenatal delivery of a therapeutic antisense oligonucleotide achieves broad biodistribution in the brain and ameliorates Angelman syndrome phenotype in mice. Mol. Ther. 32, 935–951 (2024).

29. Cohen, J. L. et al. In Utero Enzyme-Replacement Therapy for Infantile-Onset Pompe’s Disease. N. Engl. J. Med. 387, 2150–2158 (2022).

30. Shore, A. N. et al. Reduced GABAergic Neuron Excitability, Altered Synaptic Connectivity, and Seizures in a KCNT1 Gain-of-Function Mouse Model of Childhood Epilepsy. Cell Rep. 33, 108303 (2020).

31. Schwindt, P. C., Spain, W.J. Crill, W.E. Long-lasting reduction of excitability by a sodium-dependent potassium current in cat neocortical neurons. J. Neurophysiol. 61, 233–244 (1989).

32. Kim, U. Mccormick, D.A. Functional and Ionic Properties of a Slow Afterhyperpolarization in Ferret Perigeniculate Neurons In Vitro. J. Neurophysiol. 80, 1222–1235 (1998).

33. Liu, X. Leung, L.S. Sodium-activated potassium conductance participates in the depolarizing afterpotential following a single action potential in rat hippocampal CA1 pyramidal cells. Brain Res. 1023, 185–192 (2004).

34. MacLean, J. N., Zhang, Y., Johnson, B.R. Harris-Warrick, R.M. Activity-Independent Homeostasis in Rhythmically Active Neurons. Neuron 37, 109–120 (2003).

35. Marder, E. Goaillard, J.-M. Variability, compensation and homeostasis in neuron and network function. Nat. Rev. Neurosci. 7, 563–574 (2006).

36. Li, B. et al. Neuronal Inactivity Co-opts LTP Machinery to Drive Potassium Channel Splicing and Homeostatic Spike Widening. Cell 181, 1547-1565.e15 (2020).

37. Zhang, Y. et al. Rapid single-step induction of functional neurons from human pluripotent stem cells. Neuron 78, 785–798 (2013).

38. Yuan, T. et al. Coupling of Slack and NaV1.6 sensitizes Slack to quinidine blockade and guides anti-seizure strategy development. eLife 12, (2024).

39. Wang, H. et al. Slack (KCNT1) potassium channels regulate levels of proteins of the inner mitochondrial membrane. Biophys. J. 123, 250a (2024).

40. Ryner, R. F. et al. Cortical Parvalbumin-Positive Interneuron Development and Function Are Altered in the APC Conditional Knockout Mouse Model of Infantile and Epileptic Spasms Syndrome. J. Neurosci. 43, 1422–1440 (2023).

41. Gertler, T. S., Cherian, S., DeKeyser, J.-M., Kearney, J.A. George, A.L. KNa1.1 gain-of-function preferentially dampens excitability of murine parvalbumin-positive interneurons. Neurobiol. Dis. 168, 105713 (2022).

42. Shore, A. N. et al. Heterozygous expression of a Kcnt1 gain-of-function variant has differential effects on SST- and PV-expressing cortical GABAergic neurons. (2024) doi:10.7554/elife.92915.1.

43. Burbano, L. E. et al. Antisense oligonucleotide therapy for KCNT1 encephalopathy. JCI Insight 7, e146090 (2022).

44. Pantazis, C. B. et al. A reference human induced pluripotent stem cell line for large-scale collaborative studies. Cell Stem Cell 29, 1685-1702.e22 (2022).

45. Smith, R. S. et al. Sodium Channel SCN3A (NaV1.3) Regulation of Human Cerebral Cortical Folding and Oral Motor Development. Neuron (2018) doi:10.1016/j.neuron.2018.07.052.

