## Supplemental Results for "Gene therapy for targeting a prenatally enriched potassium channel associated with severe childhood epilepsy and premature death"

### Supplemental Figures

**Figure S1**
KCNT1-p.R474H neurons display large afterhyperpolarizations (AHPs) sensitive to $Ca^{++}$ activated BK and SK channel blockers

**Figure S2**
ASO-treated patient cells exhibit improved spiking properties

**Figure S3**
Prenatal emergence of Slack currents in mid-gestation in primary human neurons

**Figure S4**
ASO-treated fetal human neurons are sensitive to KCNT1 knockdown and quinidine treatment of neurons

### Supplemental Tables

**Table S1**
Electrophysiology analysis and statistics for ASO treated KCNT1 p.R474H neurons

**Table S2**
Electrophysiology values and statistics for excitability and slow AHP of ASO treated of KCNT1-p.R474H neurons

**Table S3**
Electrophysiology analysis and statistics of $K_{Na}$ primary fetal neurons

**Table S4**
Electrophysiology values and statistics for primary neuron ASO knockdown

### Supplemental Figures

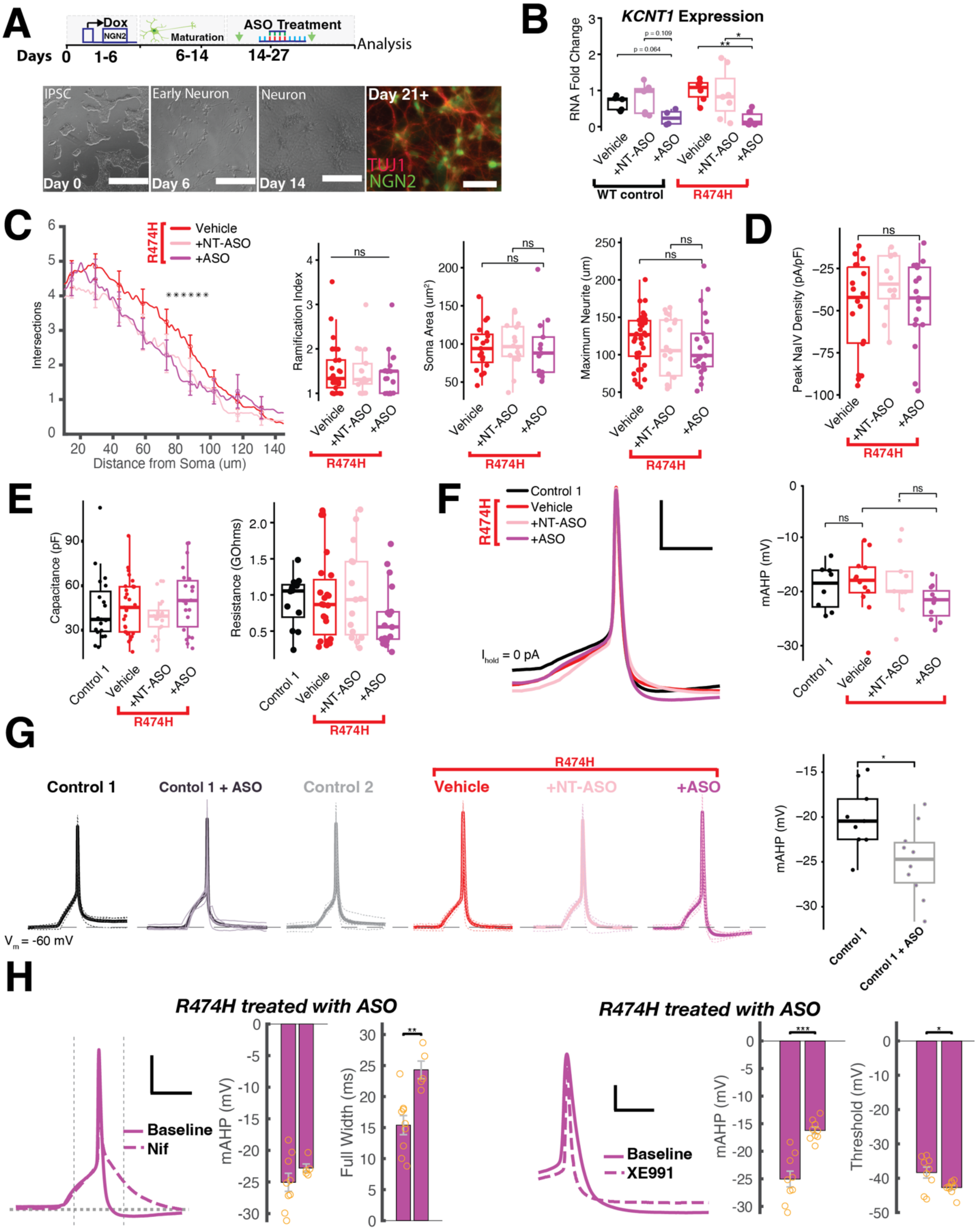

**Figure S1. KCNT1 p.R474H neurons display large afterhyperpolarizations (AHPs) sensitive to $Ca^{++}$ activated BK and SK channel blockers**

**A.** *Top*, Schematic of NGN2 differentiation of iPSCs to neurons from an individual with *KCNT1*-p.R474H variant**;** Day-0 iPSCs, Day 0-6 doxycycline treatment, Day-14 ASO treatment begins, Day 28 patch-clamp assays. *Bottom*, representative images at key steps in NGN2-directed differentiation protocol, and *Right*, immunofluorescence images of NGN2 labeling and neuronal specific marker Microtubule Associated Protein 2 (MAP2). Scale bars Day-0, 750 um; Days 6-14, 150 um; Day-21 50um. **B.** Quantitative RT-PCR analyses of *KCNT1* in control line PGP1 and p.R474H NGN2 neurons following treatment with Vehicle, ASO, or NT-ASO. Fold changes are presented with Log2 scale, and gene expression levels in vehicle-treated cells were adjusted to 1, one-sample t-test and Wilcoxon test, p-values. **C.** *Left*, Sholl analysis of neurons at Days 26-28 ($p < 0.05$ for distances between 80-100 um from cell soma). *Right*, corresponding ramification index, soma surface area, and longest primary neurite length. No significant differences observed in ramification index, soma area, and longest primary neurite length. **D.** Capacitance-adjusted values for peak inward sodium current reveals that differences in sodium influx was not the mechanism causing reduced $K^+$ current in ASO-treated cells. **E.** Analysis of capacitance and input resistance of neurons during patch-clamp analysis, no significant differences in capacitance or resistance ($p > 0.88$, $p > 0.8$, respectively). **F.** *Left*, Overlayed average current clamp recordings of a minimal stimulation resulting in a first action potential for each condition, with neurons recorded at resting potential ($I_{hold}$ = 0pA). mAHP is increased in p.R474H neurons treated with ASO, compared to vehicle ($p = 0.032$). **G.** *Left*, Representative current clamp recordings of a minimal stimulation first action potential for each condition and overlayed average, (including second control line KOLF, and control line PGP1-NGN2 neurons treated with ASO). Dashed line indicates resting potential held at -60 mV. *Right*, analysis of mAHP of PGP1-NGN2 neurons treated with ASO shows that ASO treatment enhanced mAHP in this control line similar to the effect observed in the patient line. **H.** *Left*, Average action potential of ASO-treated patient R474H neurons following bath perfusion with calcium channel blocker nifedipine (Nif, 100 uM, dashed) shows disrupted AHP kinetics compared to baseline (solid line), including a significantly greater full width ($p = 0.0025$), however mAHP was able to return to normal levels in the 200ms time window following AP peak. *Right*, Average action potential of ASO-treated p.R474H neurons with bath perfused KCNQ antagonist XE991 (100 uM, dashed) compared to baseline (solid line). *Right*, XE991 perfusion results in a significant reduction of mAHP and hyperpolarizes the AP threshold ($p < 0.001$, $p = 0.032$. *Note that reduction in mAHP is partly due to the hyperpolarized threshold from which the mAHP is measured. See Table S2 for complete values.

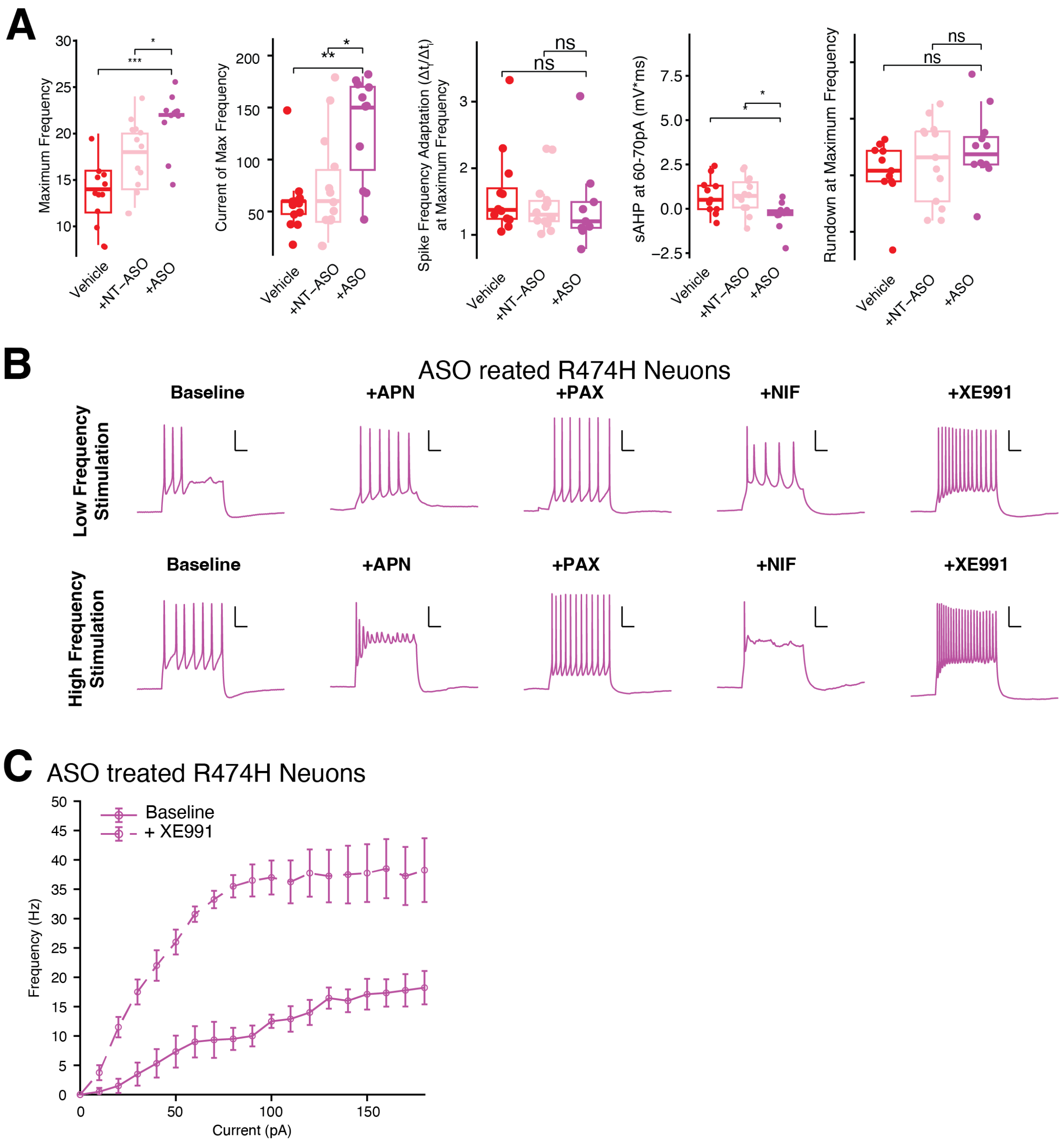

A
Maximum Frequency
Current of Max Frequency
Spike Frequency Adaptation (Δt/Δt) at Maximum Frequency
sAHP at 60-70pA (mV*ms)
Rundown at Maximum Frequency
Vehicle
+NT-ASO
+ASO
ns
B
ASO reated R474H Neuons
Baseline
+APN
+PAX
+NIF
+XE991
Low Frequency Stimulation
High Frequency Stimulation
C
ASO treated R474H Neuons
Baseline
+ XE991
Frequency (Hz)
Current (pA)

**Figure S2. ASO-treated KCNT1-p.R474H NGN2 neurons exhibit improved dynamic range of firing sensitive to SK channel blocker Apamin.**

**A.** Analysis of action potential firing properties of KCNT1-p.R474H NGN2 neurons treated with either vehicle, NT-ASO, or ASO at $V_m$=60mV with depolarizing steps (10pA steps, ranging from -10pA to 180pA). **B.** Representative spiking profiles at low and high frequency stimulation in ASO treated KCNT1-p.R474H neurons with select pharmacology, Paxalline (500nM), Apamin (50nM), Nifedipine (100 uM), and XE991 (100 uM). **C**. Input output excitability curve of ASO treated KCNT1-p.R474H neurons perfused with KCNQ antagonist XE991 showing an increased excitation across low and high input levels. $p < 0.05$ at 10 pA and 170-180 pA, $p < 0.001$ from 20-110 pA, $p < 0.01$ from 120-160 pA. Data are presented as mean ± SEM.

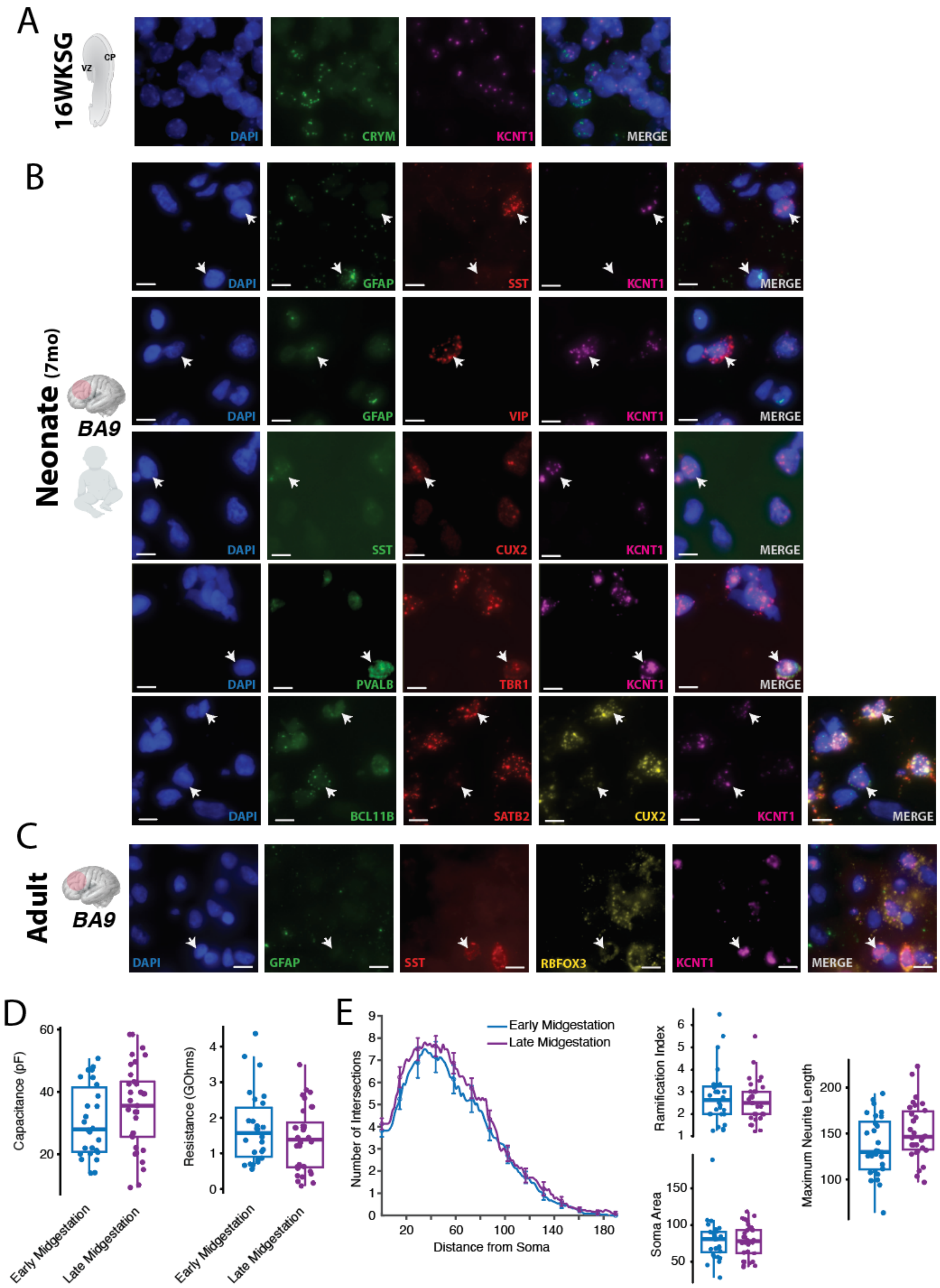
A
16WKSG
VZ
CP
DAPI
CRYM
KCNT1
MERGE
B
Neonate (7mo)
BA9
GFAP
SST
VIP
CUX2
PVALB
TBR1
BCL11B
SATB2
C
Adult
RBFOX3
D
Capacitance (pF)
Resistance (GOhms)
Early Midgestation
Late Midgestation
E
Number of Intersections
Distance from Soma
Ramification Index
Soma Area
Maximum Neurite Length

**Figure S3. Prenatal emergence of Slack currents in mid-gestation in primary human neurons**

**A.** *KCNT1* RNA *in situ* hybridization of a 16 weeks gestation (WKSG) coronal brain section from the perisylvian region demonstrate enrichment to the cortical plate (CP) and subplate marker CRYM. CP, cortical plate; VZ, ventricular zone. **B.** Analysis of *KCNT1* and cell type markers in the neonatal neocortex (7 month old), including pyramidal neuron markers TBR1, an interneuron markers (PV, VIP, and SST) and glia marker (GFAP). **C.** Analysis of KCNT1 and cell type markers in adult neocortex (Brodmann area, BA9), with cell type specific markers, including neuron markers RBFOX3, interneuron marker (SST), and glia marker (GFAP). **D.** Analysis of capacitance and input resistance of early and late midgestation primary neurons, no significant differences observed ($p = 0.208$, $p = 0.244$, respectively). **E.** *Left*, Sholl analysis of primary neurons at early and late midgestational timepoints and *Right*, corresponding ramification index, soma surface area, and longest primary neurite length. No significant differences detected in complexity, ramification index, soma area, and longest primary neurite length, suggesting cell morphology did not change while in culture (See table S3 for values).

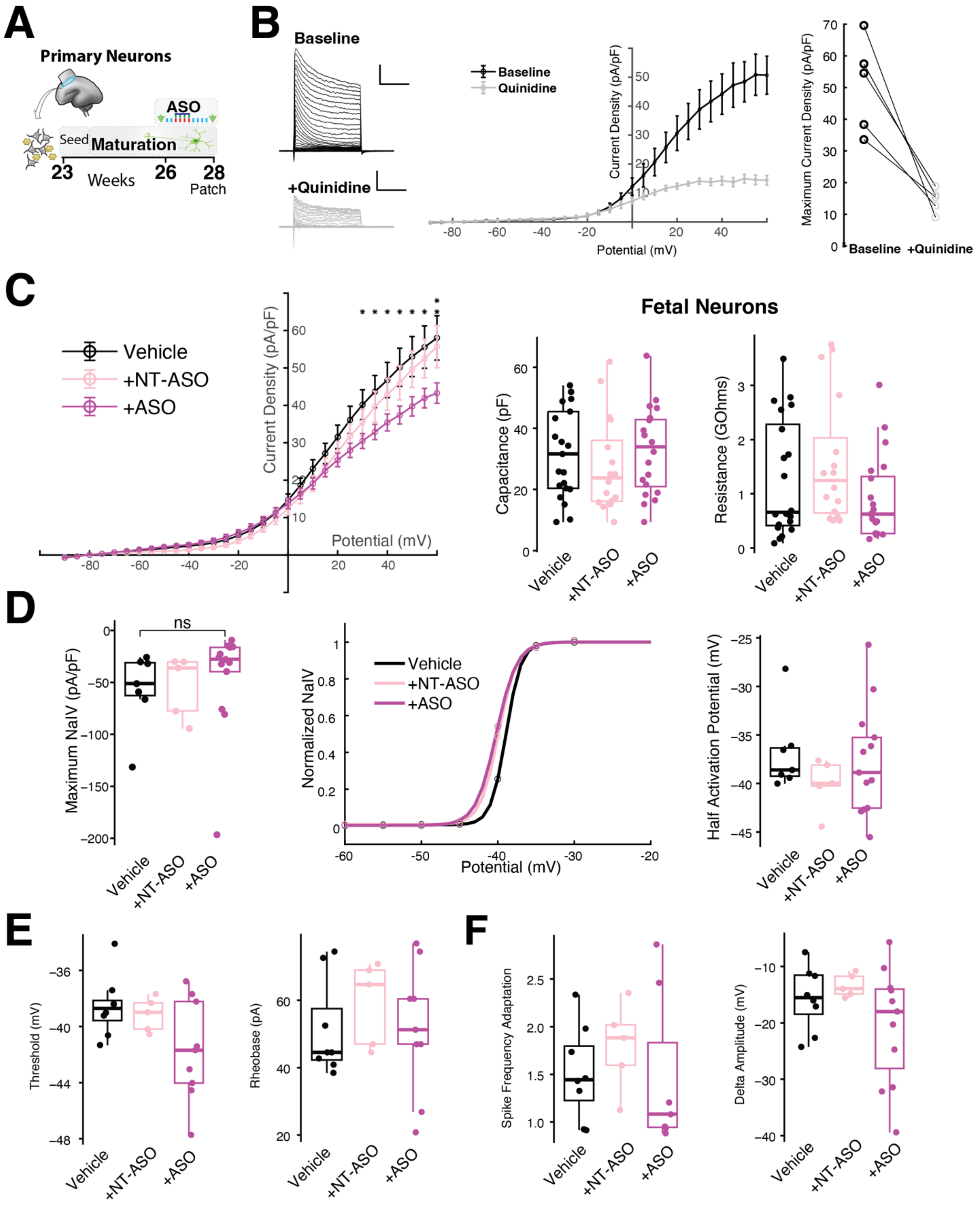


**Figure S4. ASO knockdown of $K_{Na}1.1$ (*KCNT1*) in midgestation primary human neurons, and quinidine treatment of neurons**

**A.** *Top*, Schematic of primary neuron isolation from midgestation human cortex and maturation timeline with ASO treatment**. B.** Primary fetal neuron steady-state $K^+$ current can be significantly reduced using non-selective EIMFS drug Quinidine at 100 uM ($p = 0.008$). **C.** Primary fetal neurons at 28 PCW equivalent isolated from a 23 PCW sample exhibit outward $K^+$ current densities sensitive to ASO knockdown (10uM, 14 days). ASO treatment did not significantly alter fetal neuron capacitance or input resistance during patch experiments. **D.** ASO treatment did not affect maximum $Na^+$ current magnitude or half activation potential in fetal neurons (see table S4). **E. & F.** Primary fetal neurons have comparable average thresholds, rheobase, spike frequency adaptation, and rundown across all conditions (n.s., see table S4).

| | Vehicle | +NT-ASO | +ASO | p-value | Figure |
|---|---|---|---|---|---|
| Observations | $N = 28$ | $N = 18$ | $N = 24$ | | |
| Capacitance (pF) | $46.19 \pm 3.41$ | $39.67 \pm 2.88$ | $50.61 \pm 4.59$ | $p = 0.565$ | S2-E |
| Resistance (GΩ) | $0.96 \pm 0.13$ | $1.01 \pm 0.16$ | $0.69 \pm 0.11$ | $p = 0.218$ | S2-E |
| Max $I_K$ (pA/pF) | $54.43 \pm 3.70$ | $56.35 \pm 4.14$ | $37.45 \pm 3.49$ | $p = 0.002$ ** | 2-B |
| Max $I_{Na}$ (pA/pF) | $-48.61 \pm 6.61$ | $-34.46 \pm 5.14$ | $-45.46 \pm 6.03$ | $p = 0.988$ | S2-D |

| | Vehicle | +NT-ASO | +ASO | p-value | Figure |
|---|---|---|---|---|---|
| Observations | $N = 6$ | $N = 7$ | $N = 6$ | | |
| Fold Change ($\log_2$) | N/A | $0.91 \pm 0.27$ | $0.22 \pm 0.09$ | $p = 0.013$, $p = 0.044$ */* | S2-B |

| | Vehicle | +NT-ASO | +ASO | p-value | Figure |
|---|---|---|---|---|---|
| Observations | $N = 8$ | $N = 8$ | $N = 9$ | | |
| Rheobase (pA) | $28.32 \pm 4.02$ | $31.73 \pm 4.80$ | $46.04 \pm 5.66$ | $p = 0.025$ * | 2-D |
| Threshold (mV) | $-38.76 \pm 0.91$ | $-40.36 \pm 1.11$ | $-38.35 \pm 1.60$ | $p = 0.831$ | 2-D |
| Amplitude (mV) | $64.72 \pm 3.15$ | $60.90 \pm 4.14$ | $59.93 \pm 2.92$ | $p = 0.281$ | S2-G |
| mAHP (mV) | $-18.19 \pm 0.94$ | $-18.90 \pm 0.85$ | $-25.05 \pm 1.43$ | $p = 0.001$ ** | 2-D |
| Full Width (ms) | $13.24 \pm 1.06$ | $12.84 \pm 1.05$ | $15.40 \pm 1.54$ | $p = 0.279$ | S2-G |
| Latency (ms) | $39.36 \pm 1.96$ | $35.83 \pm 3.63$ | $34.99 \pm 2.77$ | $p = 0.227$ | N/A |
| mAHP at RMP (mV) | $-18.38 \pm 1.57$ | $-18.33 \pm 2.06$ | $-22.03 \pm 0.96$ | $p = 0.032$ * | S2-F |

| | Vehicle | +NT-ASO | +ASO | p-value | Figure |
|---|---|---|---|---|---|
| Observations | $N = 32$ | $N = 18$ | $N = 21$ | | |
| Ramification Index | $1.50 \pm 0.10$ | $1.50 \pm 0.12$ | $1.45 \pm 0.10$ | $p = 0.873$ | S2-C |
| Soma Area ($\mu m^2$) | $95.62 \pm 6.74$ | $98.21 \pm 7.46$ | $94.00 \pm 11.26$ | $p = 0.596$ | S2-C |
| Max Neurite ($\mu m$) | $121.49 \pm 6.07$ | $109.70 \pm 9.07$ | $112.75 \pm 9.39$ | $p = 0.274$ | S2-C |

| Vehicle | ACSF | +PAX | +APN | p-value | Figure |
|---|---|---|---|---|---|
| Observations | $N = 8$ | $N = 11$ | $N = 10$ | | |
| Threshold (mV) | $-38.76 \pm 0.91$ | $-40.02 \pm 1.03$ | $-38.62 \pm 0.77$ | $p = 0.351$, $p = 0.965$ | 2-E |
| mAHP (mV) | $-18.19 \pm 0.94$ | $-18.62 \pm 0.87$ | $-19.21 \pm 1.10$ | $p = 0.778$, $p = 0.515$ | 2-E |
| **+NT-ASO** | **ACSF** | **+PAX** | **+APN** | **p-value** | **Figure** |
| Observations | $N = 8$ | $N = 9$ | $N = 8$ | | |
| Threshold (mV) | $-40.36 \pm 1.11$ | $-39.32 \pm 0.90$ | $-38.06 \pm 0.53$ | $p = 0.541$, $p = 0.161$ | 2-E |
| mAHP (mV) | $-18.90 \pm 0.85$ | $-18.14 \pm 1.27$ | $-18.81 \pm 1.45$ | $p = 0.423$, $p = 0.645$ | 2-E |
| **+ASO** | **ACSF** | **+PAX** | **+APN** | **p-value** | **Figure** |
| Observations | $N = 9$ | $N = 13$ | $N = 12$ | | |
| Threshold (mV) | $-38.35 \pm 1.60$ | $-39.52 \pm 0.56$ | $-39.64 \pm 1.03$ | $p = 0.262$, $p = 0.422$ | 2-E |
| mAHP (mV) | $-25.05 \pm 1.43$ | $-20.36 \pm 0.71$ | $-18.11 \pm 1.13$ | $p = 0.022$, $p = 0.005$ */** | 2-E |

| +ASO | ACSF | +NIF | +XE991 | p-value | Figure |
|---|---|---|---|---|---|
| Observations | $N = 9$ | $N = 5$ | $N = 9$ | | |
| Threshold (mV) | $-38.35 \pm 1.60$ | $-39.28 \pm 1.03$ | $-42.72 \pm 0.65$ | $p = 0.438$, $p = 0.032$ ns/* | S2-H |
| mAHP (mV) | $-25.05 \pm 1.43$ | $-22.79 \pm 0.63$ | $-16.22 \pm 0.61$ | $p = 0.298$, $p < 0.001$ ns/*** | S2-H |

**Table S1:** Electrophysiology values and statistics for KCNT1-p.R474H neurons

| | Vehicle | +NT-ASO | +ASO | p-value | Figure |
|---|---|---|---|---|---|
| Observations | $N = 11$ | $N = 13$ | $N = 12$ | | |
| Frequency at 180 pA (Hz) | $5.00 \pm 0.60$ | $7.60 \pm 2.93$ | $18.22 \pm 2.17$ | $p = 0.006$ ** | 3-C |
| Maximum Frequency (Hz) | $14.36 \pm 0.80$ | $17.23 \pm 0.98$ | $20.33 \pm 1.32$ | $p = 0.003$ ** | S3-A |
| Max Stim Current (pA) | $59.09 \pm 9.86$ | $73.85 \pm 14.12$ | $126.67 \pm 14.84$ | $p = 0.001$ ** | S3-A |
| $\Delta$Amplitude (mV) | $-9.22 \pm 1.08$ | $-7.21 \pm 1.45$ | $-4.25 \pm 1.41$ | $p = 0.011$ * | 3-D |
| Spk Freq Adaptation | $1.45 \pm 0.07$ | $1.35 \pm 0.09$ | $1.33 \pm 0.10$ | $p = 0.235$ | 3-D |
| sAHP (6-8 Hz) (mV·s) | $1.45 \pm 0.28$ | $1.18 \pm 0.13$ | $0.10 \pm 0.23$ | $p = 0.002$ ** | 3-E |
| sAHP (10-14 Hz) (mV·s) | $1.05 \pm 0.24$ | $0.52 \pm 0.19$ | $-0.01 \pm 0.20$ | $p < 0.001$ *** | 3-E |
| sAHP (16-18 Hz) (mV·s) | $0.46 \pm 0.26$ | $0.08 \pm 0.17$ | $-0.51 \pm 0.11$ | $p = 0.002$ ** | 3-E |

| Vehicle | ACSF | +PAX | +APN | p-value | Figure |
|---|---|---|---|---|---|
| Observations | $N = 11$ | $N = 8$ | $N = 11$ | | |
| Frequency at 180 pA (Hz) | $5.00 \pm 0.60$ | $22.57 \pm 4.54$ | $3.82 \pm 1.63$ | $p = 0.026$, $p = 0.682$ */ns | 3-F |

| +NT-ASO | ACSF | +PAX | +APN | p-value | Figure |
|---|---|---|---|---|---|
| Observations | $N = 13$ | $N = 6$ | $N = 7$ | | |
| Frequency at 180 pA (Hz) | $7.60 \pm 2.15$ | $21.33 \pm 7.96$ | $4.86 \pm 2.26$ | $p = 0.022$, $p = 0.446$ */ns | 3-F |

| +ASO | ACSF | +PAX | +APN | p-value | Figure |
|---|---|---|---|---|---|
| Observations | $N = 12$ | $N = 10$ | $N = 10$ | | |
| Frequency at 180 pA (Hz) | $18.22 \pm 2.17$ | $25.00 \pm 5.17$ | $3.00 \pm 0.45$ | $p = 0.271$, $p < 0.001$ ns/*** | 3-F |

| +ASO | ACSF | +NIF | +XE991 | p-value | Figure |
|---|---|---|---|---|---|
| Observations | $N = 12$ | $N = 5$ | $N = 8$ | | |
| Frequency at 180 pA (Hz) | $18.22 \pm 2.17$ | N/A | $38.25 \pm 7.44$ | N/A, $p = 0.017$ * | S3-C |

**Table S2: Electrophysiology spiking various experimental conditions of KCNT1 p.R474H neurons (data corresponds to Figure 2 and S2)**

| Fetal Neurons | Early Midgestation | Late Midgestation | p-value | Figure |
|---|---|---|---|---|
| Observations | $N = 26$ | $N = 27$ | | |
| Capacitance (pF) | $30.58 \pm 2.28$ | $33.78 \pm 2.71$ | $p = 0.208$ | S1-D |
| Resistance (GΩ) | $1.73 \pm 0.20$ | $1.36 \pm 0.19$ | $p = 0.244$ | S1-D |
| Max $I_K$ (pA/pF) | $49.97 \pm 4.12$ | $53.03 \pm 5.35$ | $p = 0.886$ | 1-D |
| Max $I_{K_{Na}}$ (pA/pF) | $23.00 \pm 7.27$ | $14.15 \pm 6.48$ | $p = 0.409$ | 1-D |
| $I_{K_{Na}}$(0 mV) (pA/pF) | $4.15 \pm 1.06$ | $11.35 \pm 1.99$ | $p = 0.003$ ** | 1-D |
| $V_{1/2}$ of $I_{K_{Na}}$ (mV) | $10.03 \pm 1.70$ | $0.39 \pm 1.14$ | N/A | 1-D |

| | Early Midgestation | Late Midgestation | p-value | Figure |
|---|---|---|---|---|
| Observations | $N = 28$ | $N = 30$ | | |
| Ramification Index | $2.79 \pm 0.24$ | $2.59 \pm 0.17$ | $p = 0.634$ | S1-E |
| Soma Area ($\mu m^2$) | $79.47 \pm 5.43$ | $78.12 \pm 4.10$ | $p = 1.000$ | S1-E |
| Max Neurite ($\mu$m) | $136.12 \pm 6.12$ | $152.37 \pm 5.58$ | $p = 0.063$ | S1-E |

**Table S3:** Electrophysiology analysis and statistics of $K_{Na}$ primary fetal neurons, **(data corresponds to Figure 3 and S3)**

| Fetal Neurons | Vehicle | +NT-ASO | +ASO | p-value | Figure |
|---|---|---|---|---|---|
| Observations | $N = 21$ | $N = 16$ | $N = 20$ | | |
| Capacitance (pF) | $31.97 \pm 3.20$ | $27.89 \pm 3.89$ | $32.63 \pm 3.15$ | $p = 0.990$ | 4-C |
| Resistance (G$\Omega$) | $1.28 \pm 0.24$ | $1.60 \pm 0.30$ | $0.90 \pm 0.17$ | $p = 0.409$ | XXX |
| Max $I_K$ (pA/pF) | $58.02 \pm 5.93$ | $55.69 \pm 5.61$ | $43.28 \pm 2.72$ | $p = 0.032$ | 4-C |
| Max $I_A$ (pA/pF) | $XXX \pm XXX$ | $XXX \pm XXX$ | $XXX \pm XXX$ | $p = XXX$ | |

| Fetal Neurons | Vehicle | +NT-ASO | +ASO | p-value | Figure |
|---|---|---|---|---|---|
| Observations | $N = 30$ | $N = 17$ | $N = 22$ | | |
| Ramification Index | $2.59 \pm 0.17$ | $2.42 \pm 0.21$ | $2.54 \pm 0.24$ | $p = 0.669$ | 4-B |
| Soma Area ($\mu m^2$) | $78.12 \pm 4.10$ | $76.66 \pm 7.34$ | $80.26 \pm 6.05$ | $p = 0.920$ | 4-B |
| Max Neurite ($\mu$m) | $152.37 \pm 5.58$ | $127.83 \pm 10.11$ | $123.79 \pm 8.79$ | $p = 0.050$, $p = 0.027$ */* | 4-B |

| Quinidine | Baseline | +Quinidine | p-value | Figure |
|---|---|---|---|---|
| Observations | $N = 5$ | $N = 5$ | | |
| Capacitance (pF) | $27.22 \pm 5.08$ | $27.04 \pm 5.55$ | $p = 0.910$ | XXX |
| Resistance (G$\Omega$) | $2.55 \pm 0.66$ | $2.90 \pm 1.05$ | $p = 0.624$ | XXX |
| Max $I_K$ (pA/pF) | $50.71 \pm 6.58$ | $14.41 \pm 1.67$ | $p = 0.008$ ** | 4-D |

**Table S4: Electrophysiology values and statistics for primary neuron ASO knockdown**
